# An innate immune circuit involving the lectin pathway, complement and lung epithelial cells enforce barrier immunity to inhaled fungal disease

**DOI:** 10.64898/2026.06.15.732416

**Authors:** Darin L. Wiesner, Gaurav Bairwa, Prashant R. Desai, Amy P. Hsu, Alex J. Whitehead, Xin He, Cleison Ledesma Taira, Lucas Dos Santos Dias, Jack A. Bibby, Claudia Kemper, Eric Karlins, Jingwen Gu, Justin Lack, Paul J. Brennan, Jigar V. Desai, Rebecca A. Brockman-Schneider, James E. Gern, George R. Thompson, Hrishikesh Kulkarni, Marcel Wüthrich, Gregory A. Demopulos, Jatin M. Vyas, Bruce Klein

**Affiliations:** Center for Immunity and Inflammation, New Jersey Medical School, Rutgers-The State University of New Jersey, Newark, NJ; Department of Medicine, New Jersey Medical School, Rutgers-The State University of New Jersey, Newark, NJ; Department of Pediatrics, University of Wisconsin School of Medicine and Public Health, Madison, WI; Laboratory of Clinical Immunology and Microbiology, National Institute of Allergy and Infectious Diseases, National Institutes of Health, Bethesda, MD 20892; Draper, Cambridge, MA; National Heart, Lung, and Blood Institute (NHLBI), National Institutes of Health (NIH), Complement and Inflammation Research Section (CIRS), Bethesda, MD 20892; Bioinformatics and Computational Biosciences Branch, National Institute of Allergy and Infectious Diseases, National Institutes of Health, Bethesda, MD; Center for Discovery and Innovation, Hackensack Meridian Health, Nutley, NJ 07110; Department of Medicine, University of California Davis Medical Center, Sacramento, CA; Divisions of Pulmonary and Critical Care and Sleep Medicine, Department of Medicine, University of California, Los Angeles, David Geffen School of Medicine, Los Angeles, CA; Omeros Corporation, Seattle, WA; Division of Infectious Diseases, Department of Medicine, Massachusetts General Hospital, Boston, MA; Department of Medicine, University of Wisconsin School of Medicine and Public Health, Madison, WI; Department of Medical Microbiology and Immunology, University of Wisconsin School of Medicine and Public Health, Madison, WI

**Keywords:** Lung bronchioles, Fungi, *Coccidioides*, Neutrophils, Complement, Lectin, Club Cells

## Abstract

The lung mucosal barrier thwarts many inhaled pathogens, including *Coccidioides*, the causative agent of Valley Fever. Because the earliest stages of pathogen recognition in the lung remain obscure, we investigated the initial events of barrier immunity in a murine model involving inhalation of *Coccidioides sp.* arthroconidia. Neutrophils accumulated rapidly, within 24 hours, near fungal spores in the bronchiolar airways. This response was driven by pattern recognition via the lectin complement pathway. Bronchiolar club cells propagated C3a signals and amplified the response via convergent C3aR and P2X7 signaling. We identified several *MBL2* and *P2RX7* polymorphisms that correlated with progressive disease in humans. Our assays revealed that these mutations caused functional impairments in C3a generation and P2X7 responsiveness. Our findings establish how complement signaling and epithelial sensing coordinate early immune responses to fungal infection, offering insights into essential host defense mechanisms and risk factors for disease progression in coccidioidomycosis.

## INTRODUCTION

Coccidioidomycosis, also known as Valley Fever, is a potentially severe respiratory and disseminated disease caused by inhalation of arthroconidia from *Coccidioides immitis* or *Coccidioides posadasii*. The spores are commonly found in the soil of arid regions of the southwestern United States through Mexico and parts of Central and South America^1^. Similar to *Histoplasma* and *Blastomyces*, *Coccidioides* is a pathogenic dimorphic fungus that transitions from a spore form in the environment to a parasitic form (i.e. spherule) in lungs of mammalian hosts. Tens of thousands of cases are reported annually, but the true incidence of infection is likely far higher due to underdiagnosis and underreporting^2,3^. Most exposed individuals clear the infection without exhibiting any obvious symptoms. However, disease occurs in a subset of about a third of infections that range from self-limiting pulmonary illness to life-threatening disseminated disease in 1 to 2 percent of infected individuals^4^.

Severe and disseminated coccidioidomycosis is influenced in part by host genetic factors. Some cases have been linked to Mendelian inherited traits impacting cells of the immune system. Such traits include mutations in the IL-12/IFN-γ pathway (*IFNGR1* and *IL12RB1*) and STAT1^5^; a gain-of-function mutation in STAT1 (e.g. *STAT1)* that dysregulates signaling in the IFN-γ/IL-12 pathway^6^; and dectin-1 pathway variants (e.g. *CLEC7A* and a *PLCG2* variants) and mutations in DUOX1 or DUOXA1 that promote hydrogen peroxide production^7^. With exception to the well-established Y238X mutation in *CLEC7A*^8^, most other mendelian mutations that correlate with fungal disease risk are relatively rare and do not account for most instances of disseminated coccidioidomycosis^9^.

The most prominent virulence trait of *Coccidioides* is the formation of a mature spherule^10^. *Coccidioides* arthroconidia inhaled into the lungs transform into titanic (20-200 μM), thick-walled spherules in response to host stressors. Hundreds of endospores fill the spherule until the spherule ruptures and releases the endospores to propagate the infection. This short delay in morphologic transition creates a window of opportunity for the host to efficiently control the infection^11^. Therefore, we reason that innate barrier immunity represents an unexplored, important checkpoint that can stall fungal growth and efficiently contain the infection.

The lung epithelium is the first host surface encountered by inhaled pathogens, where it serves not only as a physical barrier but also as a dynamic regulator of immunity^12,13^. Studies of barrier immunity in other murine models of fungal pneumonia follow a consistent, bi-phasic paradigm. In the early phase, epithelial IL-1R/MyD88 signaling induces CXCL1, CXCL5, and CCL2 to recruit neutrophils and monocytes^14,15^, while epithelial-derived GM-CSF enhances neutrophil fungicidal activity^16,17^. In the subsequent phase, CARD9-dependent pathways in hematopoietic cells, including neutrophils, further boost antimicrobial functions as innate responses transition toward lymphocyte-dependent mechanisms^15,16^. Despite these insights, the initiating signals and pathways used by epithelial cells to drive robust neutrophil recruitment to the conducting airways are poorly understood for any fungal pathogen, much less *Coccidioides*^18^.

Herein, we report a prominent role for early sensing of inhaled *Coccidioides* arthroconidia by components of the lectin-dependent complement system. We find that processed complement fragments, and particularly the anaphylatoxin C3a signal, were generated and sensed via club cell surface C3aR1 and P2RX7 receptors. The resulting downstream intracellular signaling pathways converged to prompt release of soluble signals that recruited neutrophils to the airways to restrain infection. We also report that genetic polymorphisms in several components of this early warning system render human hosts susceptible to severe and disseminated coccidioidomycosis, potentially underpinning long-standing clinical observations.

## RESULTS

### Modeling pulmonary coccidioidomycosis in mice

The relatively high exposure rates in endemic regions and broad range of disease outcomes suggest that host-pathogen interactions within an infected individual influence disease severity^18^. *Coccidioides* spores are most vulnerable to host defense before transitioning to spherules. Therefore, the initial immune responses are thought to be major determinants of disease^11^. For this reason, we focused on identifying the earliest protective immune responses at the respiratory mucosa within 24–48 hours of inhaling *Coccidioides* arthroconidia.

Due to limited treatment options, poorly defined host determinants of severe disease, and its capacity to cause disease in immunocompetent individuals, *Coccidioides* is classified as a Biosafety Level 3 (BSL-3) pathogen^19^. To enable initial, detailed studies under BSL-2 conditions, we used an attenuated strain of *C. posadasii* carrying a deletion of the *CPS1* gene (*Δcps1*)^20^. This mutation impairs spherule maturation in vitro and in vivo^20^. Notably, mature spherules typically emerge in the lungs of mice >3 days post-infection (after the endpoint of most of our experiments). Importantly, we also validated key observations with spores from a wild type, fully virulent strain of *C. posadasii* to ensure the rigor and relevance of our findings.

### Neutrophils accumulate in the bronchioles within hours of *Coccidioides* spore inhalation

To identify immune cells that interact directly with *Coccidioides* spores early in infection, mice were infected intranasally with 5 × 10⁴ CFSE-labeled *Δcps1* spores. At 24 hours post-infection, mice were injected intravenously with fluorescently conjugated anti-CD45 antibodies to distinguish circulating leukocytes from those residing in lung tissue. Lungs were then harvested, enzymatically digested, and leukocytes isolated for flow cytometric analysis. Both alveolar macrophages and neutrophils colocalized with CFSE-labeled spores, with neutrophils representing a greater proportion and absolute number of spore-associated cells (**Fig. 1A**).

**Figure 1.**
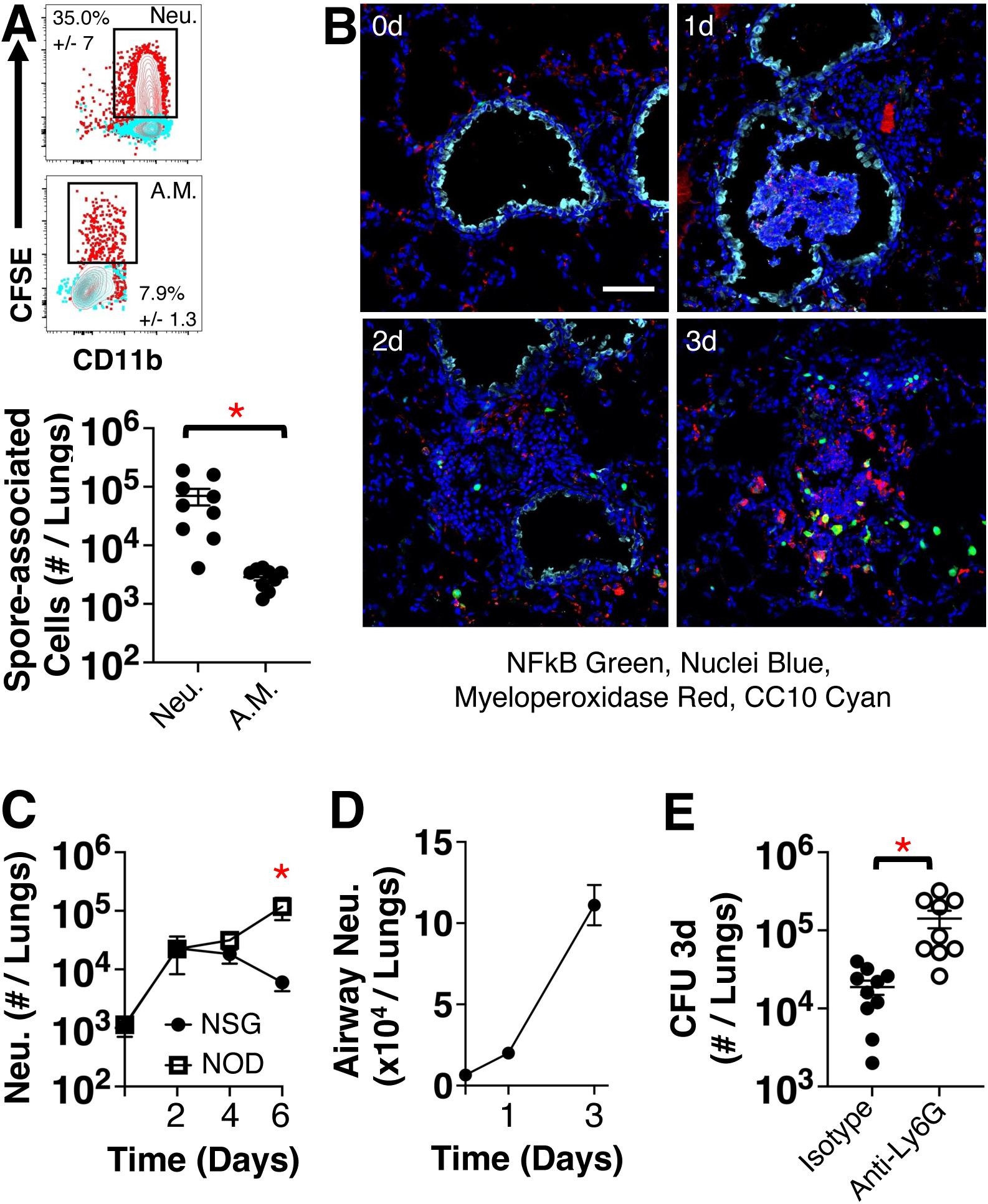
Rapid Influx of Neutrophils into the Bronchiolar Region of the Lung Following Inhalation of *Coccidioides* spores. **(A)** Representative flow plot (left) of neutrophil (top) or alveolar macrophage (bottom) association with carboxyfluorescein succinimidyl ester (CFSE)-labeled (Blue) and unlabeled (Red) spores 24 hours post infection with *Coccidioides* dCps1. Quantification (Right) of CFSE+ neutrophils or CFSE+ alveolar macrophages. **(B)** Representative confocal images from naïve and infected transgenic NF-κB-reporter mice. NFkB reporter = green, blue = DAPI/nuclei, Red = Myeloperoxidase (i.e. neutrophils) Cyan = Club Cell 10 kDa protein. Scale bar = 50μm. **(C)** Neutrophils from whole lungs of lymphocyte-deficient mice (NSG) and wild type controls (NOD). **(D)** Neutrophils bronchoalveolar lavage fluid of mice infected with fully virulent *Coccidioides.* **(E)** Colony forming units (CFU) from lungs of 3 days after mice treated with anti-Ly6G antibody/isotype control and infected with wild type virulent *Coccidioides*. All data are representative of at least 2 independent experiments. Groups indicated as “⎴” are compared by Mann-Whitney *U*. * = *p-*value < 0.05. Error bars are standard error of the mean.

To characterize the spatial and temporal dynamics of the early immune response, we infected mice and analyzed lung tissue using confocal fluorescence microscopy. At 24 hours post-infection, clusters of myeloperoxidase-positive (MPO⁺) neutrophils were observed within bronchioles lined by CC10⁺ club cells (**Fig. 1B**). By two days post-infection, this neutrophilic inflammation had shifted from the airway lumen to the peribronchiolar adventitial space within the bronchiolar bundle (**Fig. 1B**). The integrity of the CC10⁺ bronchiolar epithelium progressively eroded during this period, with complete loss by day three (**Fig. 1B**).

We observed similar patterns of neutrophil accumulation in both lymphocyte-deficient NOD-scid-gamma (NSG) mice and genetically matched NOD wild type controls (**Fig. 1C**). Lung neutrophil numbers began to diverge between these groups between days 4 and 6 post-infection, suggesting that early neutrophil recruitment progresses independently of innate lymphocyte-dependent signals.

Finally, we examined the extent and impact of neutrophil recruitment in wild-type mice infected with spores from the wild type virulent *C. posadassi* strain. Neutrophils steadily accumulated in the airways from 0 to 3 days post-infection (**Fig. 1D**), and neutrophil depletion significantly increased fungal burden in the lungs by day 3 (**Fig. 1E**). Thus, neutrophils play a requisite role in the early restraint of pulmonary infection following inhalation of *Coccidioides* arthroconidia

Taken together, these findings demonstrate a rapid, lymphocyte–independent accumulation of neutrophils in the bronchiolar region of the lung during early infection. Moreover, this rapid neutrophil response provides frontline defense, limiting the early growth of virulent *Coccidioides* in the lungs.

### The DECTIN-1/CARD9 signaling axis is dispensable for initial recognition of Coccidioides

We next investigated how the immune system recognizes inhaled *Coccidioides* spores and initiates the rapid influx of neutrophils. We hypothesized that this recognition might be driven either by dynamic changes associated with spore germination or by inherent features of the spore surface, such as microbial-associated molecular patterns. To test this, we compared neutrophil responses in mice infected with viable spores versus formalin-killed spores (FKS). Within 24 hours post-infection, both the degree of neutrophil accumulation in the airways and the bronchiolar localization of neutrophils were comparable between the two groups (**Fig. 2A, B**). These findings suggest that immune recognition is triggered by a conserved microbial pattern on the spore surface rather than a process dependent on spore germination.

**Figure 2.**
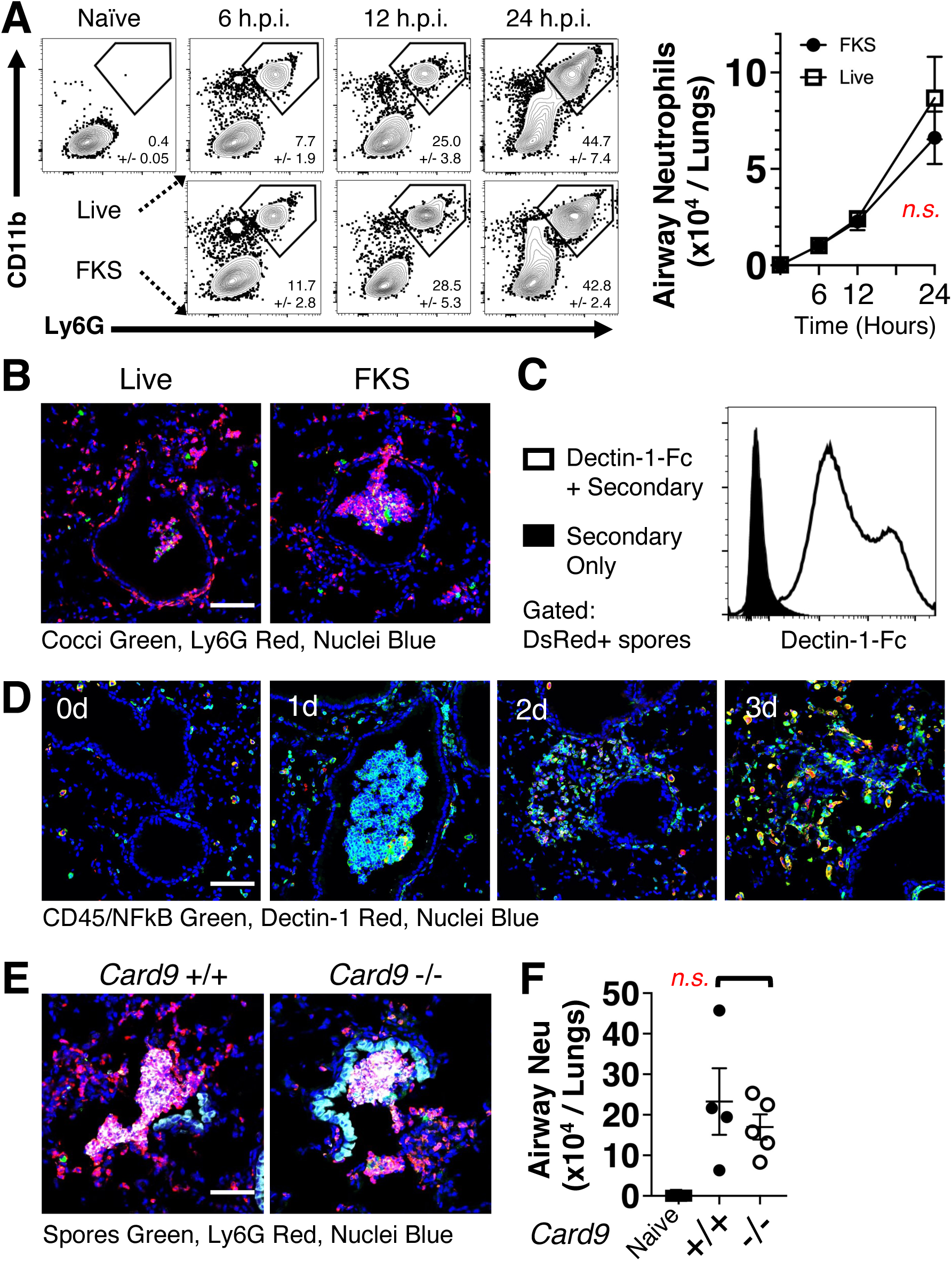
Canonical Fungal Pattern Recognition, Dectin-1-CARD9, is Dispensable for Rapid Neutrophil Recruitment. **(A)** Flow plots and graphs of airway neutrophils from lavage fluid of mice infected with *Coccidioides Δcps1*. **(B)** Representative confocal images from wild type mice 24 hours post infection with live or formalin killed spores (FKS). *Coccidioides* = green, blue = DAPI/nuclei, Red = Ly6G (i.e. neutrophils). Scale bar = 50μm. **(C)** DsRed *Coccidioides* spores analyzed for Dectin-1-Fc binding by flow cytometry. **(D)** Representative confocal images from naïve and infected wild type mice. CD45/NFkB = green, blue = DAPI/ nuclei, Red = Dectin-1. Scale bar = 50μm. **(E)** Representative confocal images from wild type or CARD9-deficient mice infected 24 hours prior. *Coccidioides* = green, blue = DAPI/nuclei, Red = Ly6G (i.e. neutrophils). Scale bar = 50μm. **(F)** Quantification of airway neutrophils of wild type and CARD9-deficient mice 24 hours post infection. All data are representative of at least 2 independent experiments. Groups indicated as “⎴” are compared by Mann-Whitney *U*. * = *p-*value < 0.05 and *n.s.* = *p-*value > 0.05. Error bars are standard error of the mean.

Next, we investigated which pattern recognition receptors (PRRs) might mediate immune detection. Dectin-1 plays a key role in sensing *Coccidioides* in mice and humans due to its expression of β-glucan in the cell wall^7,21^. We incubated DsRed⁺ spores with soluble Dectin-1-Fc, followed by fluorescently labeled anti-Fc secondary antibodies, and quantified receptor binding by flow cytometry (**Fig. 2C**). Using a similar method, we assessed binding of Dectin-2 and Mincle (**Supplemental Fig. 1A**). All three soluble receptors bound efficiently to the surface of *Coccidioides* spores. To determine whether these receptors could recognize spores in the context of host cell membrane expression, we employed cell-based reporter assays for Dectin-1, Dectin-2, and Mincle^22^. Spores of the virulent *C. posadasii* strain also activated all three receptor reporters, confirming functional recognition and signaling within the natural constraints of cell surface receptors (**Supplemental Fig. 1B**).

Although Dectin-1 is a prototypical fungal PRR and regulates myeloid cell sensing of *Coccidioides* and patient susceptibility^7^, its expression was not detectable on bronchiolar epithelial cells or the majority of neutrophils at baseline or 24 hours post-infection. Notably, Dectin-1 expression was not detected on epithelial cells across the infection, and it was evident on CD45+ leukocytes by day two and later on (**Fig. 2D**). Moreover, CARD9 deficiency, despite being the central signaling adaptor downstream of these PRRs, did not affect neutrophil accumulation in the bronchioles (**Fig. 2E**) or the total lung neutrophil numbers at 24 hours post-infection (**Fig. 2F**).

Together, these findings indicate that while multiple canonical fungal PRRs can bind surface ligands on *Coccidioides* spores, the early neutrophil response to pulmonary infection occurs independently of CARD9-mediated signaling.

### Lectin complement pathway mediates immune detection of inhaled *Coccidioides* spores

The speed and specificity of the immune response to inhaled *Coccidioides* spores suggested the involvement of a rapidly acting, broadly distributed soluble factor capable of amplifying an initial recognition signal. We hypothesized that a zymogen cascade (e.g. lectin complement pathway) would meet these criteria and mediate the early detection of *Coccidioides*. To test this, we investigated the association of key lectin pathway components with inhaled spores. These included the pattern recognition molecules mannose-binding lectin (Mbl2) and ficolin-1 (Fcn1); the serine proteases MASP1 and MASP2 that lead to formation of the lectin pathway C3 convertase, the principle enzyme complex activating the complement component C3; and the C3 cleavage products and opsonins iC3b and C3d. CFSE+ spores collected from the bronchoalveolar lavages of infected mice or the initial inoculum (negative control) were stained with fluorescently-labeled antibodies against each component and analyzed by flow cytometry. Spores recovered from lavages 24 hours post infection showed higher levels of MBL2, FCN1, MASP1 and MASP2, and iC3b labeling compared to spores from the inoculum, indicating active engagement of the lectin pathway during infection (**Fig. 3A**). In contrast, C3d was not detected above background levels (**Fig. 3A**). Confocal microscopy also revealed MBL2 deposition in and around infected bronchioles at 24 hours post-infection (**Fig. 3B**), supporting rapid in situ activation of the lectin pathway.

**Figure 3.**
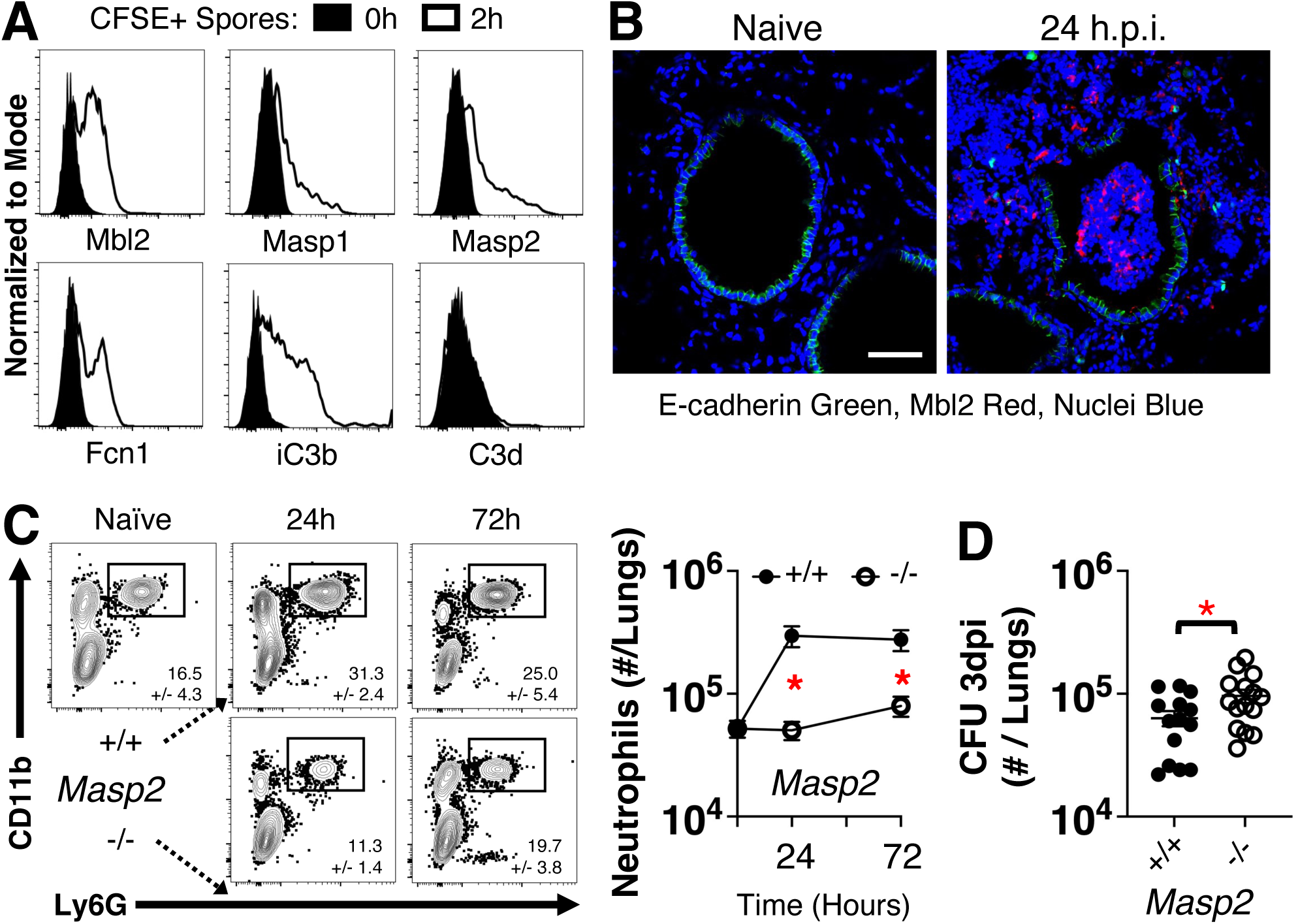
Early Recognition of *Coccidioides* Mediated by Lectin Complement Pathway. **(A)** CFSE+ *Coccidioides* spores analyzed for complement component binding by flow cytometry. **(B)** Representative confocal images from wild type mice 24 hours post infection with *Coccidioides Δcps1*. E-cadherin = green, blue = DAPI/nuclei, Red = Mbl2. Scale bar = 50μm. **(C)** Live, tissue-resident (intravascular antibody negative) neutrophils from whole lungs of wild type mice or *MASP2*^−/−^ and wild type mice infected with *Coccidioides Δcps1*. **(D)** Colony forming units (CFU) from lungs of wild type or *MASP2*^−/−^ mice and infected 3 days prior with wild type virulent *Coccidioides.* All data are representative of at least two independent experiments. Groups indicated as “⎴” are compared by Mann-Whitney *U*. * = *p-*value < 0.05. Error bars are standard error of the mean.

MASP2 is the main activator of the lectin pathway and is immediately downstream of MBL2 and FCN1. To assess the functional importance of the lectin pathway in host defense, we infected *MASP2*^−/−^ mice with attenuated *Δcps1* spores and quantified neutrophil recruitment at 0-, 24-, and 72-hours post-infection. While baseline neutrophil levels were similar between wild-type and knockout mice, *Masp2*^−/−^ mice failed to elicit an increase in lung neutrophils following infection (**Fig. 3C**). Consistent with this, intravenous treatment of wild-type mice with MASP2-blocking antibodies also significantly reduced neutrophil accumulation in response to infection (**Supplemental Fig. 2**). Finally, infection of *Masp2*^−/−^ mice with the wild type, virulent *C. posadasii* spores resulted in significantly higher lung fungal burden compared to wild-type controls at 3 days post-infection (**Fig. 3D**). Together, these findings demonstrate that the lectin complement pathway is rapidly activated upon inhalation of *Coccidioides* spores and plays a critical role in neutrophil recruitment and early fungal containment, highlighting its function as a key component of frontline pulmonary immunity.

### Complement component C3a signals via bronchiolar club cells to drive neutrophil recruitment

Complement cleavage products C3a and C5a are well-known inflammatory mediators. Within 6 hours of *Coccidioides* infection, C3a levels sharply increased in the bronchoalveolar lavage fluid of infected mice (**Fig. 4A**), whereas C5a levels showed little to no change during the first 24 hours (**Fig. 4A**). Consistent with these findings, incubating spores in fresh serum for one hour resulted in a larger increase in C3a production than C5a (**Supplemental Fig. 3**).

**Figure 4.**
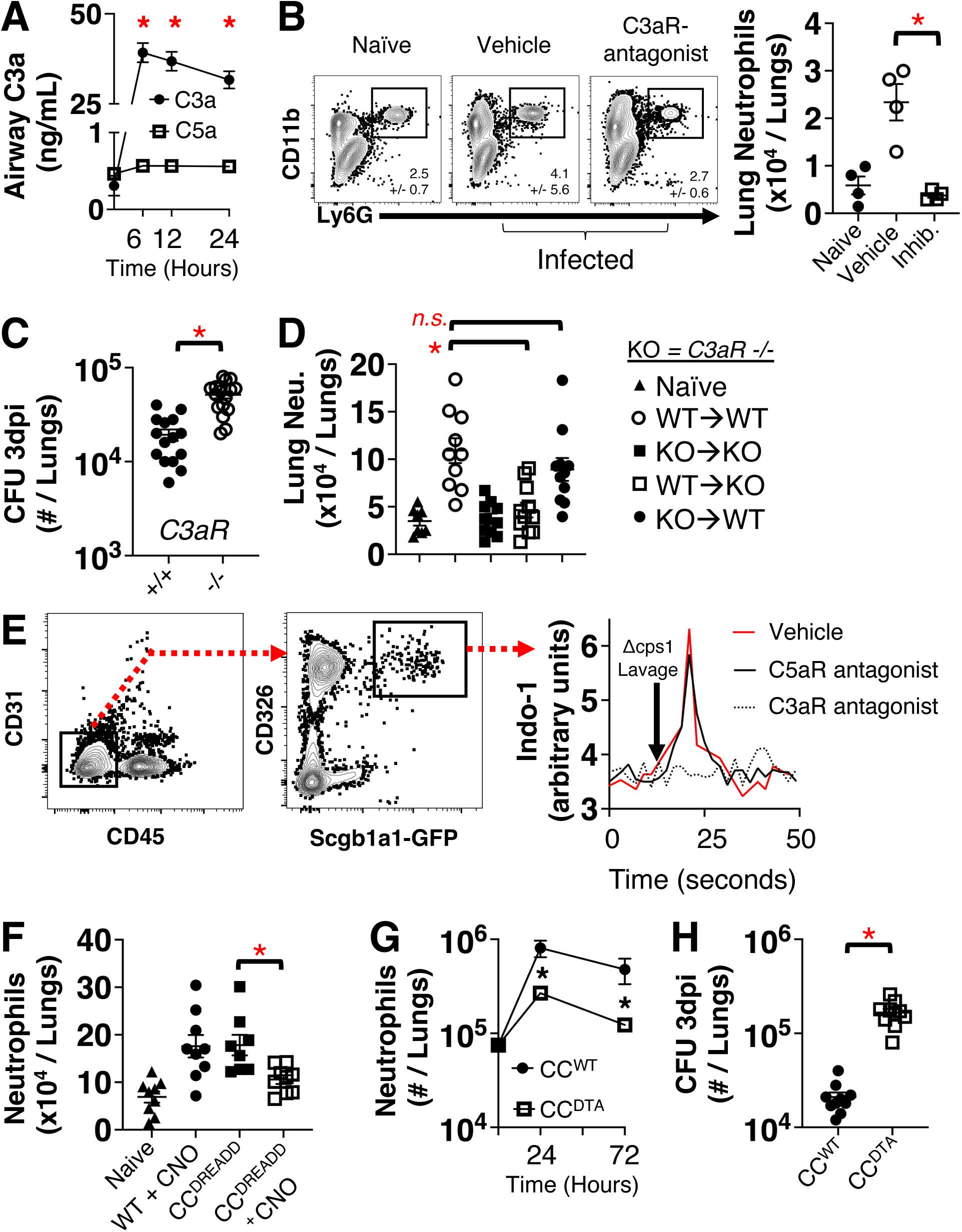
Bronchiolar Club Cells Respond to C3a and Initiate Neutrophil Recruitment. (**A**) C3a and C5a measured from lavage fluid collected from wild type mice at the indicated time points. (**B**) Live, tissue resident (intravascular antibody negative) neutrophils from whole lungs of wild type mice treated with C3a receptor antagonist (SB290157; 1.5mg/kg) or vehicle control and infected with *Coccidioides Δcps1*. (**C**) Colony forming units (CFU) from lungs of wild type or *C3AR1* −/− mice that were infected 3 days prior with wild type virulent *Coccidioides.* (**D**) Neutrophils from whole lungs of chimeric mice 3 days post infection with *Coccidioides Δcps1*. (**E**) Representative flow gating (left) and calcium flux data (right) of gated Scgb1a1+ Club Cells cells loaded with calcium indicator dye and pretreated with complement receptor antagonists (C3aR antagonist SB290157 10μM; C5aR antagonist PMX205, 500nM). Cells were briefly incubated with lavage fluid from infected mice and processed immediately by flow cytometry (**F**) Neutrophils from whole lungs of wild type and club cell-specific DREADD mice treated with CNO (GPCR inhibited) or not (wild phenotype) 3 days post infection with *Coccidioides Δcps1*. (**G**) Neutrophils from whole lungs of club cell-cre (wild phenotype) or club cell-cre x LSL-diphtheria toxin (club cell-deficient) mice infected with *Coccidioides Δcps1* (**H**) Colony forming units (CFU) from lungs of mice indicated in panel (G) that were infected 3 days prior with fully virulent *Coccidioides.* All data are representative of at least 2 independent experiments. Groups indicated as “⎴” are compared by Mann-Whitney *U*. * = *p-*value < 0.05 and *n.s.* = *p-*value > 0.05. Error bars are standard error of the mean.

To assess the role of C3a in neutrophil recruitment, we treated mice with a C3a receptor (C3aR) antagonist and observed a significant reduction in lung neutrophil accumulation 24 hours post-infection in response to *Δcps1* spores (**Fig. 4B**). Moreover, global *C3ar1*^−/−^ mice infected with virulent *C. posadalii* spores exhibited significantly higher fungal burdens in the lungs at three days post-infection compared to wild type controls (**Fig. 4C**).

To determine whether the target of C3a signaling was of hematopoietic or non-hematopoietic origin, we generated bone marrow (BM) chimera mice. We transferred wild-type BM into either wild-type or *C3ar1*⁻^/^⁻ recipients, and vice versa. Following infection with attenuated *Δcps1* spores, neutrophil responses at 72 hours post-infection were significantly elevated in both WT→WT and *C3ar1*⁻^/^⁻→WT chimeras and were not significantly different from one another (**Fig. 4D**). In contrast, neutrophil responses in both WT→*C3ar1*⁻/⁻ and *C3ar1*⁻/⁻→*C3ar1*⁻/⁻ chimeras did not rise above baseline, naïve levels (**Fig. 4D**). These results demonstrate that C3aR activation on non-hematopoietic, cells is critical for driving the early neutrophil response to *Coccidioides* infection of the lung.

Club cells are the predominant epithelial cell type in the bronchioles of mice and are thus positioned to be among the first host cells to encounter inhaled *Coccidioides* spores. To determine whether club cells respond to complement-derived inflammatory signals, we tested whether bronchoalveolar lavage fluid from infected mice elicits calcium signals through C3a or C5a receptors. GFP⁺ club cells were isolated and enriched from club cell-GFP reporter mice and loaded with a ratiometric calcium dye. Lavage fluid from infected mice induced a rapid calcium flux in the GFP⁺ club cells, a response that was abolished by C3a receptor antagonism but unaffected by C5a receptor blockade (**Fig. 4E**).

C3aR is a G protein-coupled receptor (GPCR). To establish the functional contribution of GPCR signaling in club cells during infection, we used a chemogenetic approach in CC^DREADD^ mice, in which GPCR signaling can be selectively inhibited in club cells upon treatment with clozapine-N-oxide (CNO). Wild-type mice treated with CNO and CC^DREADD^ mice without CNO both mounted normal neutrophil responses to *Coccidioides* infection (**Fig. 4F**). In contrast, CC^DREADD^ mice treated with CNO exhibited significantly impaired neutrophil recruitment (**Fig. 4F**), indicating that GPCR signaling in club cells is critical for initiating the neutrophilic response.

Local production of C3 by club cells influences club cell survival during *Pseudomonas* infection^23^. We therefore investigated the role of C3 produced by club cells in neutrophil recruitment and anti-fungal resistance. Mice with loxP sites flanking a *C3*-IRES-TdTomato transgene report C3 expression and crossing these mice with Scgb1a1-creERT2 mice allows for club cell-specific deletion of C3^24^. Cre-negative, C3-TdTomato mice infected with *Coccidioides* spores showed signs of C3-reporter expression by bronchiolar club cells, and the reporter activity was absent in Cre-positive, C3-TdTomato mice (**Supplemental Fig. 4A**). We next infected the phenotypically wild type Cre-negative mice and the Cre-positive mice (i.e. C3 conditionally deleted in club cells) with *Δcps1* Coccidioides. The neutrophil response at 24 hours was not significantly different in the two strains of mice **(Supplemental Fig. 4B).** Also, wild type and club cell, *C3*-deficient mice did not exhibit differences in CFU when infected with spores of the virulent *Coccidioides* strain. Thus, club cells produce C3 locally in this model, but local production by these cells is dispensable in neutrophil accumulation and disease control following infection.

To further investigate the role of club cells in innate immunity, we used CC^DTA^ mice, in which diphtheria toxin expression conditionally ablates club cells. These mice showed markedly reduced accumulation of lung neutrophil numbers over the first 72 hours of infection with *Δcps1* spores (**Fig. 4G**). Moreover, the mice failed to control infection with spores of the wild type, virulent strain (**Fig. 4H**). Collectively, these findings identify club cells as key mediators of C3a-induced neutrophilic inflammation and demonstrate that their GPCR signaling is essential for mounting an effective early immune response to *Coccidioides* infection. Additionally, club cells are indispensable in mounting early resistance against progressive pulmonary infection with a virulent strain of this fungus.

### C3aR signaling regulates chemotaxis and genes promoting ATP generation

To understand how club cells respond to C3aR stimulation after *Coccidioides* infection, we performed bulk RNA-sequencing of FACS sorted club cells from wild type or global *C3ar1^−/−^*mice. We plotted the top 50 differentially expressed genes (DEGs) between the groups at 0, 6 and 12 hours after infection (**Fig. 5A**). To identify gene programs uniquely engaged in the wild type club cells after infection, we performed gene ontology (GO) enrichment analysis on DEGs from the 6– and 12-hour timepoints (**Fig. 5B**).

**Figure 5.**
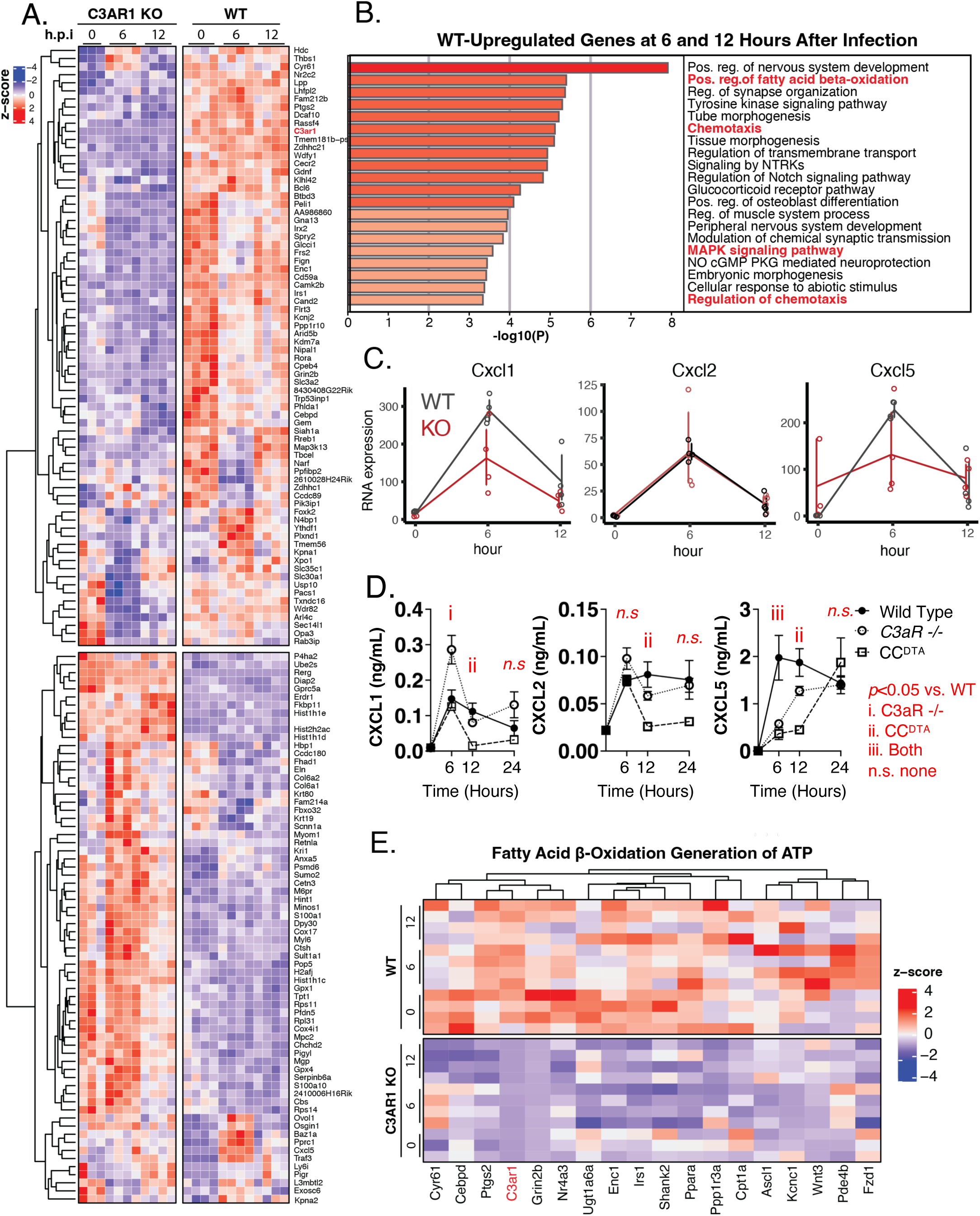
Club Cell C3aR1 regulates chemotaxis and fatty acid oxidation genes after infection. Wild type and *C3ar1^−/−^* strains of C57Bl/6 mice were infected with spores of *Δcps1* strain *of C. posadasii*. **(A)** Top 50 differentially expressed genes (DEGs) between *C3ar1*^−/−^ and wild type genotypes in club cells at baseline and at seral timepoints after infection. **(B)** Metascape Gene Ontology (GO) terms for genes upregulated (FDR<0.05, absolute LFC >0.6) in the wild type genotype after infection. **(C)** Relative mRNA expression of known neutrophil chemotaxis genes. **(D)** Quantification of chemokines in bronchoalveolar lavage fluid. **(E)** Heatmap of individual genes from the fatty acid beta oxidation GO term.

GO analysis identified differential regulation of chemokine genes known to act as neutrophil attractants (**Fig. 5C**). *Cxcl1* and *Cxcl5* mRNA production was suppressed in club cells lacking *C3ar1* receptor expression, but no difference was observed in *Cxcl2* at the transcript level. To distinguish which of these chemokines are translated and secreted in response to infection, we performed ELISAs on BAL fluid harvested 6, 12, and 24 hours after infection (**Fig. 5D**). Of the three proteins, Cxcl5 was produced in the greatest amounts, and was also suppressed in mice lacking C3aR. A similar trend was observed in Cxcl2, albeit with a smaller effect size. Cxcl1 was depressed in global club cell knockout mice, but the effect of *C3AR1* knockout was unclear.

In addition to chemotaxis, our GO analysis identified the presence of *C3AR1* as a positive regulator of 18 fatty acid β-oxidation genes (**Fig. 5E**). This metabolic process is known to generate large amounts of ATP, which can modulate cell effector function during infection^25,26^. We found that the beta oxidation genes were largely upregulated in wild type mice regardless of infection status, with the exception of *Pde4B*, *Fzd1* and Wnt3, which peaked at 6 hours post infection. Interestingly, C3ar1 was included as part of the ontology term.

### Elevation of extracellular ATP levels in the airways is dependent on C3a-C3aR axis during pulmonary *Coccidioides* infection

C3aR activation is known to influence metabolism^27^. In support of this notion, our RNA-seq results revealed *C3ar1*-dependent upregulation of genes involved in β-oxidation and ATP production (**Fig. 5E**), and C3aR signaling on macrophages has been shown to induce ATP secretion by these cells^26^. We thus reasoned that, as *Coccidioides* infection progresses, the intracellular pool of ATP may be released into the surrounding milieu, since extracellular ATP (eATP) is a well-established damage-associated molecular pattern (DAMP). Accordingly, we quantified ATP levels in bronchoalveolar lavage samples from wild type mice. Indeed, these samples showed a significant increase in eATP levels at 12– and 24-hours post-infection (**Fig. 6A**), whereas samples from *C3ar1^−/−^* mice exhibited a significant reduction in eATP levels, and those from club cell-DTA mice, even more pronounced deficits (**Fig. 6A**). Further, we treated mice with 2-deoxyglucose (2-DG), a glycomimetic that impairs glycolysis as a major source of ATP generation. Neutrophil accumulation in the lungs of 2-DG–treated mice was unaffected at 24 hours post-infection (**Fig. 6B**), indicating that early-stage ATP production may primarily rely on C3aR-mediated β-oxidation. However, as *Coccidioides* infection progressed, neutrophil recruitment was sharply impaired in 2-DG–treated mice compared to untreated or mock-treated controls at 72 hours (**Fig. 6B**). These findings suggest a biphasic shift in club cell metabolism from β-oxidation to glycolysis to sustain eATP production, which may be critical for neutrophil recruitment during early pulmonary infection with *Coccidioides*.

**Figure 6.**
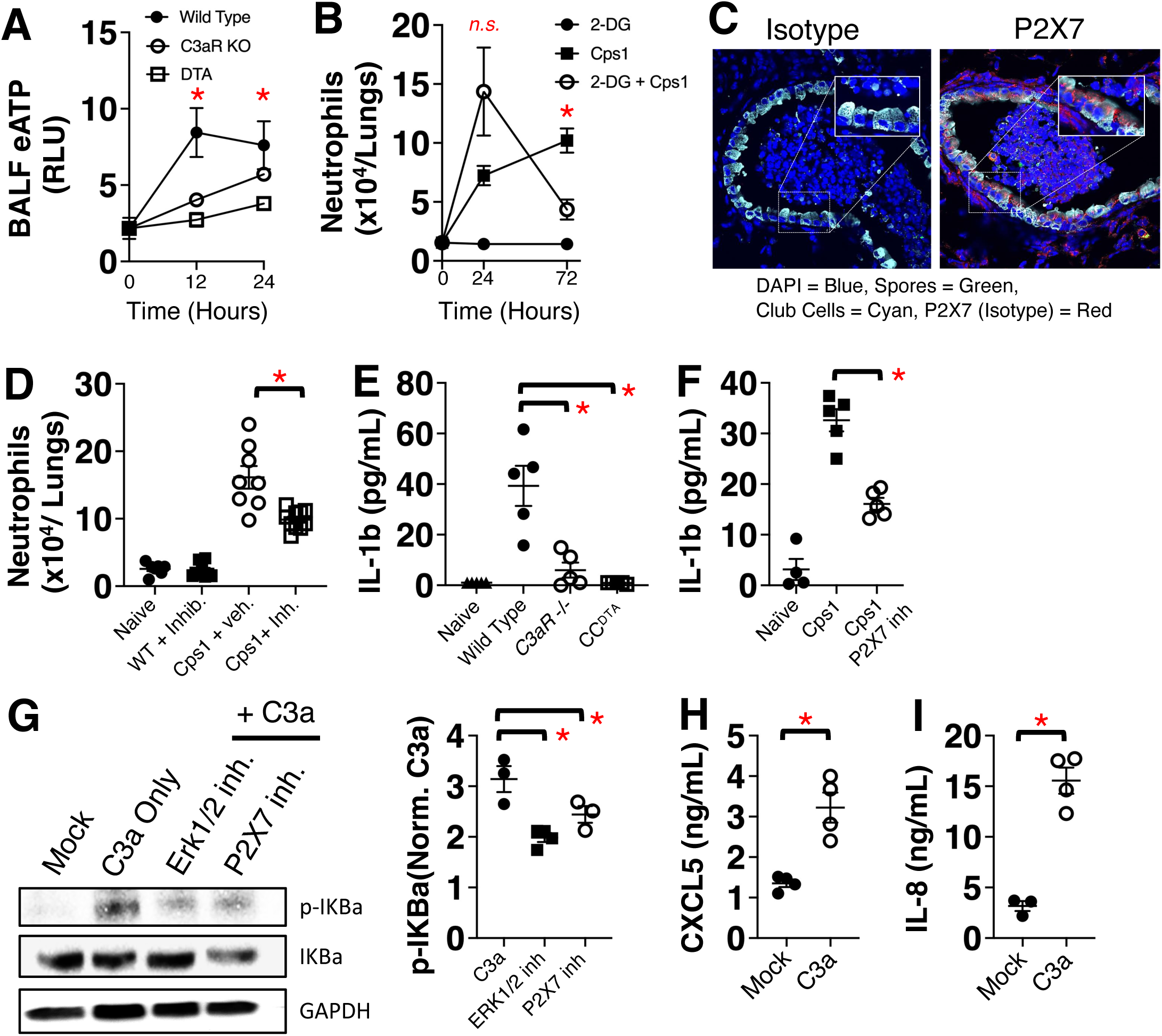
Extracellular ATP and P2X7 promote neutrophil recruitment during pulmonary *Coccidioides* infection: **(A)** Extracellular ATP (eATP) levels quantified from the BALF collected from wild type, *C3ar*1−/− or club cell DTA mice at 12h and 24h post infection. **(B)** Treatment with 2-Deoxy Glucose (i.p.1mg/kg) influences neutrophil recruitment in lungs at 72 hours postinfection in mice infected with *Δcps1 Coccidioides* spores. **(C)** Expression of P2X7 receptor on club cells in *Δcps1* infected mice at 24h postinfection. Image acquired at 60X: DAPI, blue; *Δcps1* spores-green; club cells-cyan, P2X7 or Isotype control-red. **(D)** Impact of P2X7 blockade by Az10606120 (i.p. 2mg/ kg) upon the recruitment of neutrophils in the lungs in mice infected with *Δcps1* spores. **(E)** IL-1β levels in BALF collected from *Δcps1* infected wild type, *C3ar1*−/− or club cell DTA mice. *or* **(F)** P2X7 (Az10606120) antagonist treated mice**. (G)**. SAEC were apically stimulated by C3aR agonist (100µM) for 12 hr, Erk1/2 inhibitor (20µM; FR18020) was added for 30 min prior to C3aR stimulation. P2X7 antagonist (4µM; Az10606120) was added with C3aR agonist. Densitometry quantification of p-IKBa normalized to mock-treated samples and GAPDH. SAECs were apically stimulated with C3aR agonist (100µM) for 12h and CXCL5 (**H**) was measured in apical medium. After C3aR stimulation for 24 hr, IL-8 **(I)** was quantified in the basal medium.

### eATP sensing by P2X7 favors neutrophil recruitment during pulmonary *Coccidioides* infection

eATP is a chemoattractant for neutrophils, and this DAMP is sensed by various immune and non-immune cells through purinergic receptors^28^. In this context, we observed expression of the purinergic receptor P2X7 on murine club cells (**Fig. 6C**). Importantly, P2X7 antagonist–treated mice showed a reduction in neutrophil recruitment, whereas no such effect was observed in the mock/vehicle-treated group (**Fig. 6D**). Thus, P2X7 sensing of eATP promotes neutrophil accumulation at the site of infection. P2X7 activation is known to induce release of the canonical proinflammatory cytokine IL-1β, and we found elevated levels of the product in the bronchiolar lavage fluid of wild type mice, but not *C3ar1*^−/−^ or club cell-DTA mice after *Coccidioides* infection (**Fig. 6E**); inhibition of P2X7 also impaired IL-β accumulation in the lavage fluid (**Fig. 6F**). These findings suggest a potential convergent signaling axis on club cells involving both C3aR and P2X7, similar to that observed in monocytes^26^.

### C3a stimulation of airway epithelial cells induces Erk1/2 phosphorylation and caspase-1 activation

We further investigated the C3a–C3aR interaction and its influence on intracellular and convergent signaling pathways in lung epithelial cells. Initially, we stimulated a human epithelial cell line, A549, either with C3a alone or in combination with LPS priming for 15, 30, and 60 minutes. Cellular extracts from A549 cells stimulated with C3a alone showed elevated levels of phosphorylated Erk1/2 at 30 minutes, which peaked at 60 minutes (**Supplemental Fig. 5A**). A549 cells stimulated with C3a following LPS priming exhibited greater increases in phospho-Erk1/2 levels as early as 15 minutes, with further increases at 30 and 60 minutes (**Fig. S5A**). Moreover, prolonged stimulation of A549 cells with C3a after LPS priming also resulted in higher levels of activated caspase-1 (**Supplemental Fig 5B**). Importantly, inhibition of Erk1/2 phosphorylation or antagonism of P2RX7 after stimulation with C3a each impaired caspase-1 activation (**Supplemental Fig. 5C**). Taken together, these results suggest that the C3a–C3aR axis in human lung epithelial cells promotes caspase-1 activation via Erk1/2 phosphorylation and convergent P2X7 signaling.

### SAECs respond to C3a by NF-κB activation and secretion of neutrophil recruiting chemokines

To extend findings from our mouse model and immortalized cell lines, we examined *primary* human small airway epithelial cells (SAECs) differentiated at the air–liquid interface. Surface expression of C3aR and P2X7 on club cells was confirmed by flow cytometry (**Supplemental Fig. 6A**). Unlike A549 cells, apical stimulation of SAECs with C3a did not induce caspase-1 activation, but instead triggered phosphorylation of IκBα, a factor associated with NF-κB activation (**Supplemental Fig. 5D**; and **Fig. 6G**). C3a mediated NF-κB signaling was dependent on Erk1/2 and P2X7, as inhibition of Erk1/2 phosphorylation or pharmacological antagonism of P2RX7 resulted in reduced IκBα phosphorylation (**Fig.6G**). Furthermore, C3a stimulation promoted production of the neutrophil-recruiting chemokines CXCL5 and IL-8 (**Fig. 6 H and I**). These results in human lung epithelial cells mirror findings described in blood monocytes, involving convergent signaling between C3aR and P2X7^26^.

### Variants in *MBL2* and *P2RX7* are associated with disseminated and chronic pulmonary coccidioidomycosis in patients

After identifying components of the lectin and other pathways important in the host response to *Coccidioides* infection in mice, we analyzed two cohorts of coccidioidomycosis patients with extrapulmonary disseminated coccidioidomycosis (n = 111; 31 with meningitis) or chronic pulmonary disease (n = 59)^7^. We specifically queried *MBL2*, *P2RX7*, and *C3AR1* in these cohorts. We first compared the 111 patients with disseminated disease against ancestry-matched controls from 1000 Genomes, using controls (**Fig. 7A**).^29^. A significant variant burden in *MBL2* was observed in patients compared to controls (Adj. P value, 0.03), while *P2RX7* approached significance (Adjusted [Adj.] P-value, 0.07). *C3AR1* variants were not significantly associated with disseminated disease. Examination of specific variants within the genes revealed that *MBL2 p.R52C* was significantly associated with disseminated disease in these patients*; P2RX7* p.R270C approached significance, while no variants within *C3AR1* were identified as associated with having disseminated disease.

**Figure 7.**
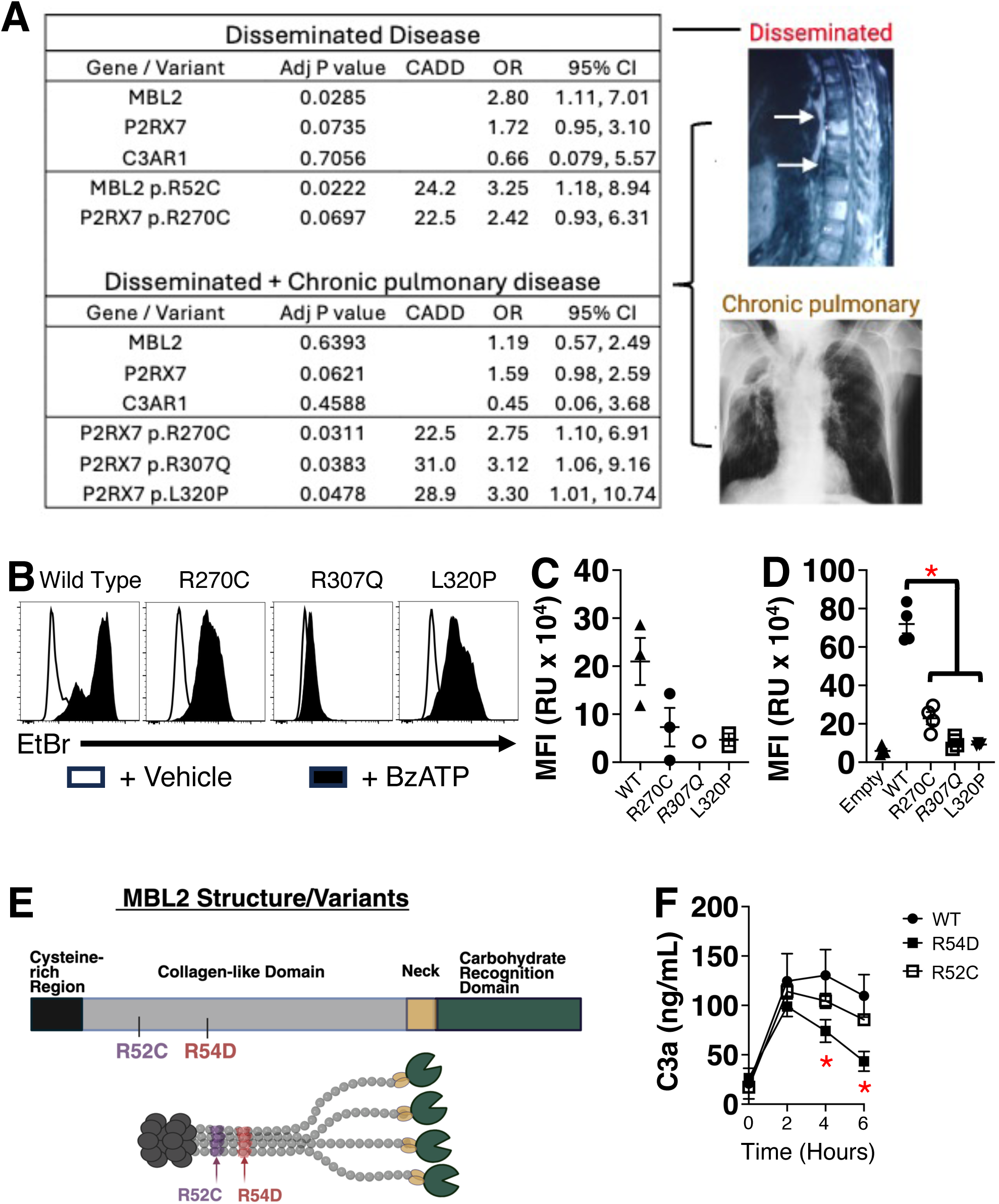
Genetic variation in and *MBL2* and *P2RX7* is associated with chronic and disseminated coccidioidomycosis in patients. (A) Variant burden analysis of disseminated (upper) or disseminated plus chronic pulmonary (lower) coccidioidomycosis patients for *MBL2*, *P2RX7*, and *C3AR1* at the gene or predicted-damaging variant level using a case-control study for risk. (B&C) *P2RX7* variants associated with risk (R270C, R307Q and L320P) were tested for response to ATP stimulation, as measured by reduced uptake of ethidium bromide. Peripheral blood mononuclear cells (PBMCs) from healthy donors and individuals harboring *P2RX7* variants were incubated with the P2X7 agonist BzATP and ethidium bromide to assess functional activity by flow cytometry. Data depicted as flow plots (**B**) and Mean Fluorescent Intensity (MFI) (C). **(D)** HEK cells transfected with wild type or variant forms of *P2RX7* tested for ethidium bromide uptake upon BzATP stimulation. **(E)** Structure and location of MBL2 variants. **(F)** Complement activation and generation of C3a upon incubation of *Coccidioides* spores with serum from individuals harboring *MBL2* variants G54D or R52C.

Since chronic pulmonary infection also is considered a severe, long-term form of coccidioidomycosis, we expanded the analysis to also include individuals with chronic pulmonary disease. We defined “chronic pulmonary coccidioidomycosis” as refractory disease lasting >12 months without extrapulmonary dissemination despite anti-fungal therapy. This second analysis included both the patients with chronic pulmonary disease (n = 59) or those with disseminated disease (n = 111)^7^. We again performed 1:4 case: control matching using 1000 Genomes controls. In this analysis, the *P2RX7* variant burden again approached significance (Adj. P-value, 0.06), while neither *MBL2* (Adj. P-value, 0.6393) nor *C3AR1* (Adj. P-value, 0.4588) were associated when chronic pulmonary and disseminated disease were combined. Examination of specific variants revealed that three *P2RX7* variants, p.R270C (Adj. P-value 0.03), p.R307Q (Adj. P-value, 0.04), and p.L320P (Adj. P-value, 0.05) are positively associated with these severe forms of coccidioidomycosis (**Fig. 7A**).

### Human variants in *P2RX7* (R270C, R307Q and L320P) result in impaired responses to eATP

In our case-control analysis, *P2RX7* variants (R270C, R307Q and L320P) were associated with elevated risk of chronic pulmonary and disseminated coccidioidomycosis. We performed functional analysis of *P2X7* variants using peripheral blood mononuclear cells from healthy human subjects. Upon sensing eATP, *P2X7* undergoes rapid conformational changes that result in increased permeability of its ligand and intracellular signaling that modulates inflammatory responses^30^. By gating on the monocytes, we tested the response to ATP stimulation as measured by uptake of ethidium bromide dye as described^31^. We tested samples displaying the wild type *P2RX7* sequence and samples with single nucleotide polymorphisms (SNPs) at R307Q, R270C or L230P (**Figs. 7B and 7C**). The R307Q SNP, a known dysfunctional variant of *P2RX7*^31^, showed dampened levels of ethidium bromide uptake compared to wild type. Importantly, R270C and L320P each showed a severe decrement in uptake of ethidium bromide dye, indicating defective responses to eATP by P2X7 by these cells (**Fig. 7C**). Additionally, we transiently transfected HEK cells with wild type *P2RX7* or the respective variants to validate our findings in primary cells. Uptake of ethidium bromide upon ATP stimulation was again sharply impaired in cells expressing the *P2RX7* variants (**Fig. 7D**). Taken together, an impairment in eATP responsiveness due to R307Q, R270C and L320P SNPs in *P2RX7* may dampen early recruitment of neutrophils and other immune cells during infection, potentially contributing to host susceptibility to severe forms of coccidioidomycosis.

### *MBL2* variants in the collagen-like domain impair sustained complement activation upon exposure to *Coccidioides* spores

Our experiments in a mouse model of pulmonary coccidioidomycosis revealed that innate immune responses to the spores require the complement pathway, particularly C3a (**Fig. 3 and 4**) generated via the lectin pathway components MBL2 and MASP2. Furthermore, our case control study of human patients demonstrated an association between *MBL2* variants (R54D and R52C) and increased risk of disseminated coccidioidomycosis (**Fig. 7A**). Both *MBL2* variants implicated in our study are located in the collagen-like domain of the protein, which is responsible for the oligomerization of Mbl2 (**Fig. 7D**)^32^. We therefore investigated the functional consequence of these variants on the generation of C3a. We identified healthy subjects displaying these variants and tested their plasma. We assessed the capacity of wild type and implicated *MBL2* variants (R54D and R52C) to activate the complement pathway in response *Coccidioides* spores *in vitro* by quantifying generation of C3a from plasma. The kinetics of C3a generation were impaired with both variants. All variants tested showed a time dependent defect in the production of C3a. The *MBL2*-R54D variant failed to sustain C3a production at the 4 h and 6 h timepoints and the MBL2-R52C variant exhibited a significant reduction in C3a levels at 6 h (**Fig. 7E**). Thus, *MBL2* variants R54D and R52C impair the ability to sustain C3a production in response to *Coccidioides* spores, suggesting that impaired complement activation in such individuals may predispose to disseminated disease.

## DISCUSSION

Our study reveals a rapid, neutrophil-dominated immune response to inhaled *Coccidioides* spores in a mouse model of coccidioidomycosis. Within 24 hours of infection, neutrophils accumulate in the bronchioles in close association with fungal spores. This early response is driven by the lectin pathway-mediated activation of complement in the lung, via MBL2 and MASP2, and resulted in the generation of the anaphylatoxin C3a. Lung club cells sense C3a via C3aR, a GPCR signaling receptor that triggers production of neutrophil chemoattractant (e.g. CXCL5) and ATP release from epithelial cells. Extracellular ATP (eATP), acting via P2X7, further amplifies neutrophil recruitment and IL-1β production in a murine model. Upon analysis of a large cohort of patients with various forms of coccidioidomycosis, we linked polymorphisms in Mbl2 and P2X7 with the risk of disseminated and chronic pulmonary forms of the disease. Finally, serum and blood samples from subjects with these polymorphisms demonstrated corresponding impairments in sustained complement activation required in innate resistance to *Coccidioides* infection.

Neutrophilic inflammation concentrated in the bronchiolar regions of the lungs of *Coccidioides* infected mice. We assume this area is where fungal spores also deposit. However, the exact location of the spores is technically challenging to determine due to delicate attachment of spores to the airways soon after infection combined with fluid displacement of fungal spores during tissue processing^33^. The conducting airways or bronchioles do not have resident immune cells as the terminal airspace has with alveolar macrophages^13^. Therefore, the host has evolved an efficient mechanism to rapidly detect an infectious threat and relay this signal with velocity and precision to coordinate a mass migration of marginated/circulating neutrophils to the exact location of infection.

Innate humoral immunity, including complement, is far-reaching and fast-acting. In particular, the lectin complement pathway comprises several classes of pattern recognition components. Mannose-binding lectin-C (Human = MBL-2), ficolin-A (Human – M-Ficolin), and pentraxin 3 are known to bind carbohydrates on the surface of *Aspergillus* and mediate protective inflammation^34–37^. Surfactants also contribute to lectin pathway activation with fungal lung infection^38^, but due to the location of the immune response in our model that occurs in the bronchioles away from the alveoli that produce surfactants, we did not prioritize investigating surfactants. Mbl-C and ficolin-A directly and pentraxin 3 indirectly converge on MBL-associated serine protease 2 (Masp2) to initiate complement enzymatic cascade and activation^39,40^.

We identified airway club epithelial cells as early sensors of *Coccidioides* spores and uncovered the C3a–C3aR axis as the driver of neutrophil recruitment in the infected airway. While complement activation in the bronchioles leads to abundant C3a generation, we found that epithelial C3aR, rather than club cell–derived C3, governs neutrophil influx during early infection. This finding contrasts with *Pseudomonas* pneumonia where club cells served as the chief source of C3a, preventing cellular damage^23^. Club cells and their C3a-C3aR axis were not only required for the early accumulation of neutrophils, but without the cells, the animals failed to contain the infection.

The epithelial response to C3a appears to diverge across species. In mice, we found IL-1β accumulation in the lung airways in the setting of convergent C3aR1–P2X7 signals, implying inflammasome activation. In human SAECs, however, C3aR signaling instead drove NF-κB activation and CXCL5 and IL-8 production. Since CXCL5 and IL-8 exert well-established roles in neutrophil chemotaxis and activation in the airway, this pathway offers a clear mechanism by which club cells mobilize neutrophils during early infection. Such epithelial-driven neutrophil recruitment may provide a critical layer of host defense by limiting spore survival and preventing transition to spherule formation within the human lung.

An inactive lectin complement pathway is the most common primary human immunodeficiency^41^. Polymorphisms in *MBL2* and *MASP2* reduce serum MBL2 concentrations and prevent proper MBL2-MASP2 complex formation^42,43^. Lectin pathway mutations have been linked with fungal infection in transplant and cancer patients^44,45^. Based on ancestry, black people are more likely to harbor mutations in *MBL2* or *MASP2,* rendering the lectin pathway inactive^46,47^. Black people are also a risk group for disseminated coccidioidomycosis^48^.

We translated our mouse model findings to the human condition. We addressed the long-standing question of host susceptibility to severe and disseminated disease. Our human genetic analyses revealed *P2RX7* and *MBL2* variants associated with increased risk of severe or disseminated coccidioidomycosis. *MBL2* polymorphisms, particularly the structural variants within exon 1 affecting arginine residues at positions 52 and 54, are known to impair the collagen-like domain required for assembly of higher-order oligomers, which represent the functionally active form of the protein^32,49^. Consistent with these results, we found that serum from individuals carrying the MBL2 R52C and R54D variants exhibited reduced generation of C3a following exposure to *Coccidioides* spores. Such a defect likely curtailed early recruitment of neutrophils during *Coccidioides* infection, impairing host defense of fungal invasion.

Extracellular ATP (eATP) in the airways acts as a canonical damage-associated molecular pattern (DAMP). eATP can be sensed by two major classes of purinergic receptors: ligand-gated ion channels of the P2X family (P2X1–P2X7) and G protein–coupled P2Y receptors (P2Y1, P2Y2, P2Y4, P2Y6, and P2Y11–P2Y14). As P2Y receptor signaling has been primarily linked to type 2 immune responses, we focused on P2X7 given its established role in driving inflammation through inflammasome activation and IL-1β production^50,51^. We detected expression of P2X7 on the surface of club cells in mice and within the bronchioles of human lung tissue. The functional relevance of genetic variation in P2X7 is underscored by the well-characterized R307Q polymorphism, which alters receptor activation^31^. In this study, we additionally identified two novel variants, R270C and L320P, that showed impaired response to eATP stimulation. Structural studies suggest that R307Q and R270C may disrupt ATP binding potential, whereas variation at L320 likely affects allosteric modulation, a mechanism coupling ligand binding and pore opening of P2X7^52,53^. Taken together, these variants may potentially weaken the process of innate immune cell recruitment and over all host defense against *Coccidioides i*nfection.

Our findings highlight the central role of the lectin complement pathway, lung epithelial cells, and neutrophils that enforce front-line immune defense in both mice and humans. The variants we identified in these pathways impair ATP sensing or sustained C3a generation when tested with functional assays. To date, a limited number of patients with disseminated disease have been investigated; they exhibited monogenic mutations associated with primary immunodeficiencies, most within the IL-12/IFN-γ and STAT3 axes^5^. A recent study uncovered damaging variants in *CLEC7A* and *PLCG2* among patients with disseminated disease that was associated with impaired production of β-glucan–stimulated TNF-α from PBMC. *DUOX1* or *DUOXA* variants that impaired H_2_O_2_ production also are overrepresented in discovery and validation cohorts in that study. While those studies have highlighted components of myeloid and lymphocyte function, our work advances the new concept that elements of the complement system in cooperation with the lung stroma and its receptors represent the first and earliest known line of defense against inhaled *Coccidioides* arthroconidia and damaging variants in pathway and receptor genes render humans vulnerable to poor outcomes.

### Limitations of the study

Our studies focused on neutrophil recruitment and accumulation as a surrogate for protective immunity. However, neutrophils require additional signals at the mucosal barrier that further augment their function and fungal killing. Because no fluorescent viability-reporter strain of *Coccidioides* is currently available, direct assessment of neutrophil-mediated killing of fungal spores in vivo was not possible. Although club cell–derived C3 production was dispensable in neutrophil recruitment, other cellular sources of C3 may compensate to maintain the local complement pool. Thus, C3 from club cells or other sources such as stromal cells may still contribute to neutrophil-dependent killing of *Coccidioides* at the airway barrier.

Mutations in Dectin-1 (*CLEC7A*) are established risk factors for fungal disease, including coccidioidomycosis^7,8^. However, we detected no baseline expression of Dectin-1 in mouse lung epithelial cells, consistent with prior work showing that adenoviral overexpression is required to elicit Dectin-1–dependent epithelial responses to *Aspergillus* infection^54^. Downstream of Dectin-1, NF-κB signaling can proceed through two distinct intermediates: CARD9, a SYK-dependent adaptor that promotes canonical NF-κB activation via RelA (p65)/p50 assembly, and Raf1, a SYK-independent pathway that modifies RelB to drive non-canonical NF-κB signaling^55^. We also did not explore an alternate signaling pathway via NFAT through PLCG2, Ca^++^ and calcineurin^7^. In our model, CARD9 was dispensable for the rapid recruitment of neutrophils to the lungs following *Coccidioides* infection. This result is consistent with prior bone marrow chimera studies demonstrating that CARD9-dependent signaling is confined to the hematopoietic compartment and becomes evident only after two or more days of infection^15,16^. Collectively, these findings suggest that Dectin-1-CARD9 signaling is unlikely to initiate the earliest innate responses to *Coccidioides* infection.

However, several important caveats temper dismissal of Dectin-1-CARD9 in our model. First, all experiments were performed in C57BL/6 mice, which harbor a mutation in exon 3 of *Clec7a* that alters the Dectin-1 stalk region and reduces surface expression and function^56,57^. Second, CARD9 captures only a subset of Dectin-1 signaling outputs, leaving alternative pathways such as Raf1^7^ intact. Because global Raf1 deficiency is embryonically lethal^58^ and a club cell-specific conditional knockout was unavailable, we were unable to directly assess epithelial Raf1-dependent signaling. Finally, our analyses focused on neutrophil recruitment rather than effector function, precluding conclusions about CARD9-dependent regulation of neutrophil antifungal activity. As a result, our model is not optimized to directly compare the relative contributions of Dectin-1-mediated pattern recognition and the lectin complement pathway, nor do these data exclude an important role for Dectin-1 in fungal sensing at later stages or in distinct cellular compartments.

We used an inoculum of 10^5^ *Δcps1* spores for infection of mice, which may be higher than the natural infectious dose; it also was several logs more than inoculum we used to infect mice with spores of the wild-type strain C735. The dose chosen for *Δcps1* spores was selected based on original mouse studies by the group that engineered this attenuated isolate^59^. Our approach allowed us to dissect the epithelial cell-driven innate mechanisms *in vivo* under controlled, reproducible conditions. Other work examining innate immune responses to *Coccidioides* has also used inocula in the range of 10⁵–10⁶ arthroconidia of an avirulent *Coccidioides* strain^60,61^.

Resource availability: Darin L. Wiesner, Bruce S. Klein, Gregory A. Demopulos

Lead contact: Darin L. Wiesner

Materials availability: Darin L. Wiesner, Bruce S. Klein

## DATA AND CODE AVAILABILITY

- RNA-seq data will be deposited at GEO with an accession number and made publicly available as of the date of publication.
- Original western blot images will be deposited at Mendeley and made publicly available as of the date of publication. Microscopy data reported in this paper will be shared by the lead contact upon request.
- Any additional information required to reanalyze the data reported in this paper is available from the lead contact upon request.

## ACKNOWLEDGEMENTS

We thank Mark Orbach and John Galgiani (University of Arizona) for providing the mutant strain *Δcps1* of *C. posadasii* and ChiungYu Hung (University of Texas, San Antonio) for providing the wild-type strain C735 of this organism.

Funded by grants from NIH including R01 AI168370 and R37AI035681 (BSK), R00 AI141622 (JVD), R01HL166449 (HK), U19 AI104317 (JEG), R01AI150181 (JMV); support from Dr. Margaret and Alan Hill, and the Burden Family Fund for Coccidioidomycosis (GRT); the Intramural Research Program of the NIH, National Institute of Allergy and Infectious Diseases (NIAID) (APH); and National Heart Lung and Blood Institute (NHLBI) Intramural Research Program grant ZIA/hl1006223 (CK).

## AUTHOR CONTRIBUTIONS

Conceptualization: DLW, GB, PD, BSK, CK, AJW, APH, JEG, GAD, MW; Methodology: JAB, CK, AJW, RAB, APH, HK, JEG, JV, GAD, LDSD; Investigation, DLW, GB, PD, JVD, JAB, JD AJW, RAB., APH, JV; GAD, GRT, EK, JG, JL, CL; Writing draft, DLW, GB, PD, APH, BSK; Writing, review and editing, all authors; Funding, BSK; Resources, GAD, HK, GRT, APH; and Supervision, DLW, BSK. CK.

## DECLARATION OF GENERATIVE AI OR AI-ASSISTED TECHNOLOGIES

During the preparation of this work, the author(s) did not use generative AI or AI-assisted technologies.

## SUPPLEMENTAL FIGURE LEGENDS

**Supplemental Figure 1.**
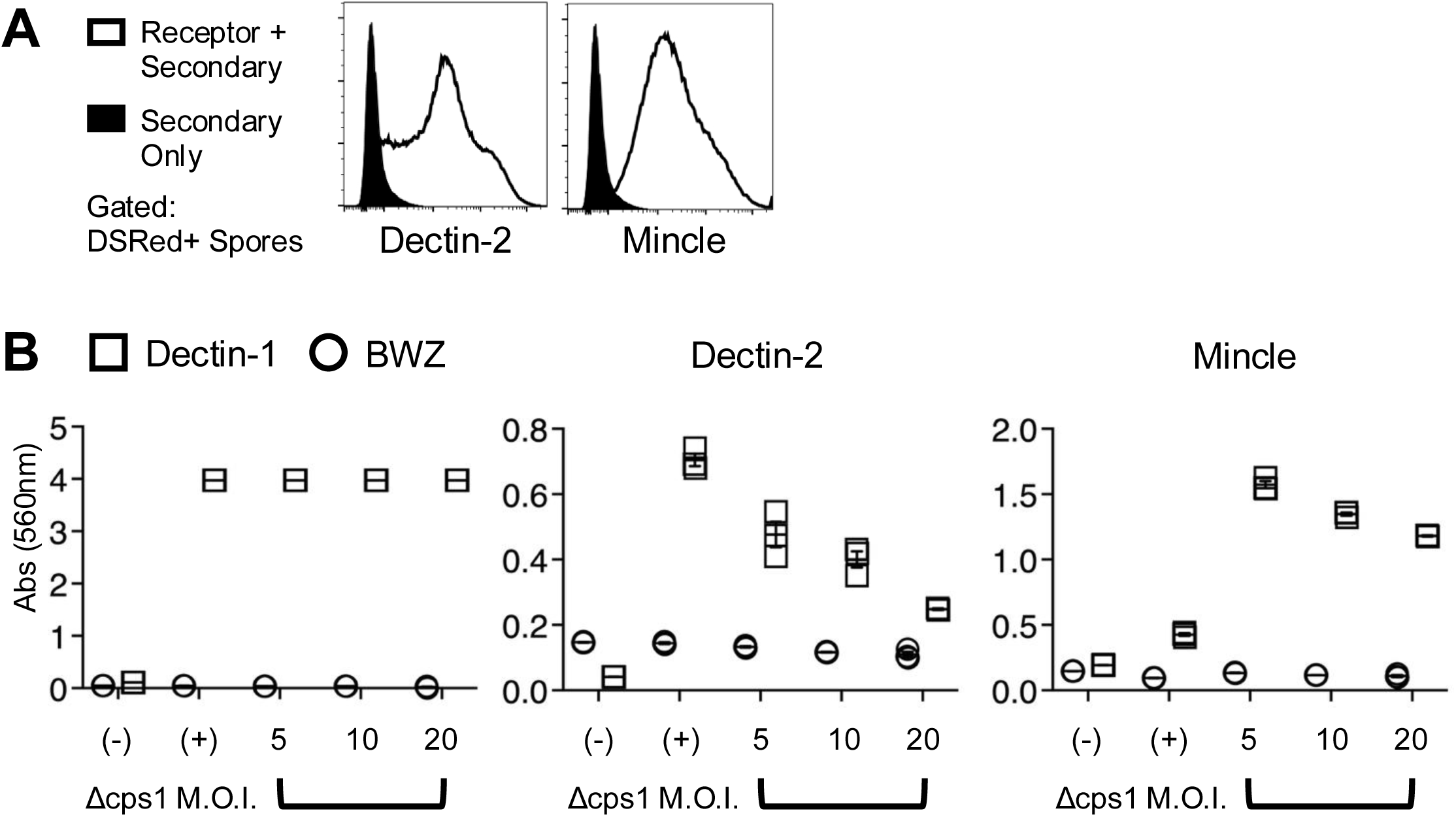
Coccidioides spores are bound by C-type lectin receptors. (**A**) DsRed⁺ spores incubated with soluble Dectin-1-Fc, followed by fluorescently labeled anti-Fc secondary antibodies, and quantified receptor binding by flow cytometry. (**B**) Cell based reporter assay or spores incubated at a range of MOI. Abs 560nm is directly proportional to signal strength. All data are representative of at least 2 independent experiments. Error bars are standard error of the mean.

**Supplemental Figure 2.**
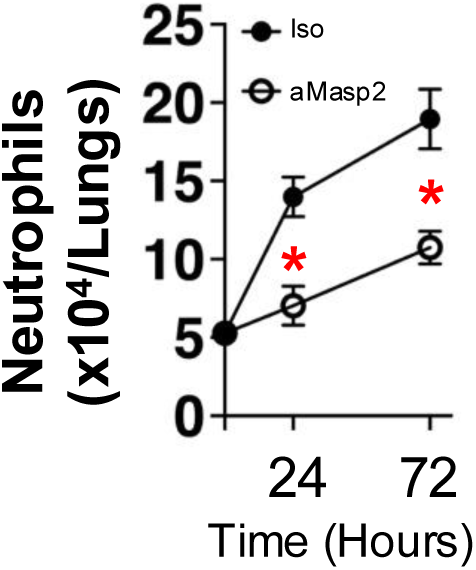
Masp2 blocking antibody inhibits neutrophil accumulation in the lungs of *Coccidioides* infected mice. Mice were treated with blocking antibody IP prior to infection. Neutrophils were quantified from lung digests. All data are representative of at least 2 independent experiments. Groups are compared by Mann-Whitney *U*. * = *p-*value < 0.05. Error bars are standard error of the mean.

**Supplemental Figure 3.**
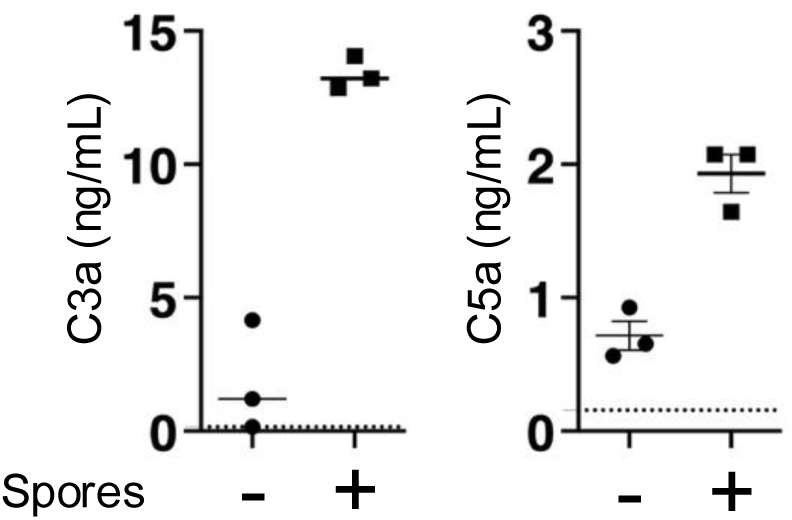
More C3a than C5a is produced when *Coccidioides* is incubated with fresh mouse serum. (**A**) *Δcps1* spores incubated with fresh mouse serum for 1 hour. C3a and C5a were quantified by ELISA. All data are representative of at least 2 independent experiments. Groups are compared by Mann-Whitney *U*. * = *p-*value < 0.05. Error bars are standard error of the mean.

**Supplemental Figure 4.**
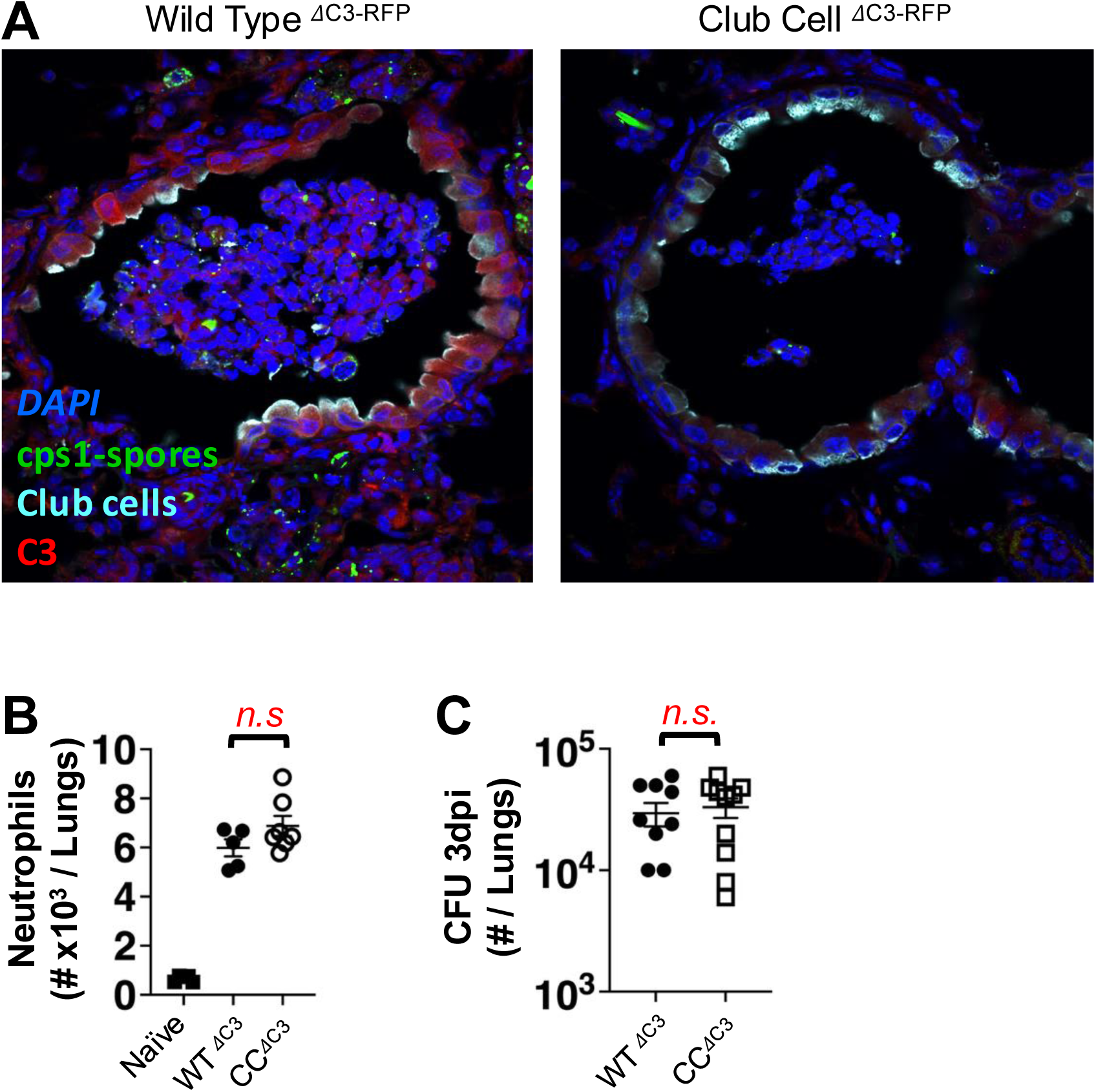
Local production of C3 by club cells does not impact immunity to *Coccidioides*. (**A**) Confocal images from wild phenotype (Scgb1a1-creER neg, C3-RFP floxed) and club cell C3-deleted mice (Scgb1a1-creER positive, C3-RFP floxed) 24 hours post infection with *Δcps1*. Blue = DAPI, Green = spores, Cyan = club cells, Red = C3. (**B**) Neutrophils from lavage of mice infected with dCps1. (**C**) Colony forming units from lungs of mice infected with the fully virulent strain of *Coccidioides*. All data are representative of at least 2 independent experiments. Groups indicated as “⎴” are compared by Mann-Whitney *U*. n.s. = *p-*value > 0.05. Error bars are standard error of the mean.

**Supplemental Fig 5.**
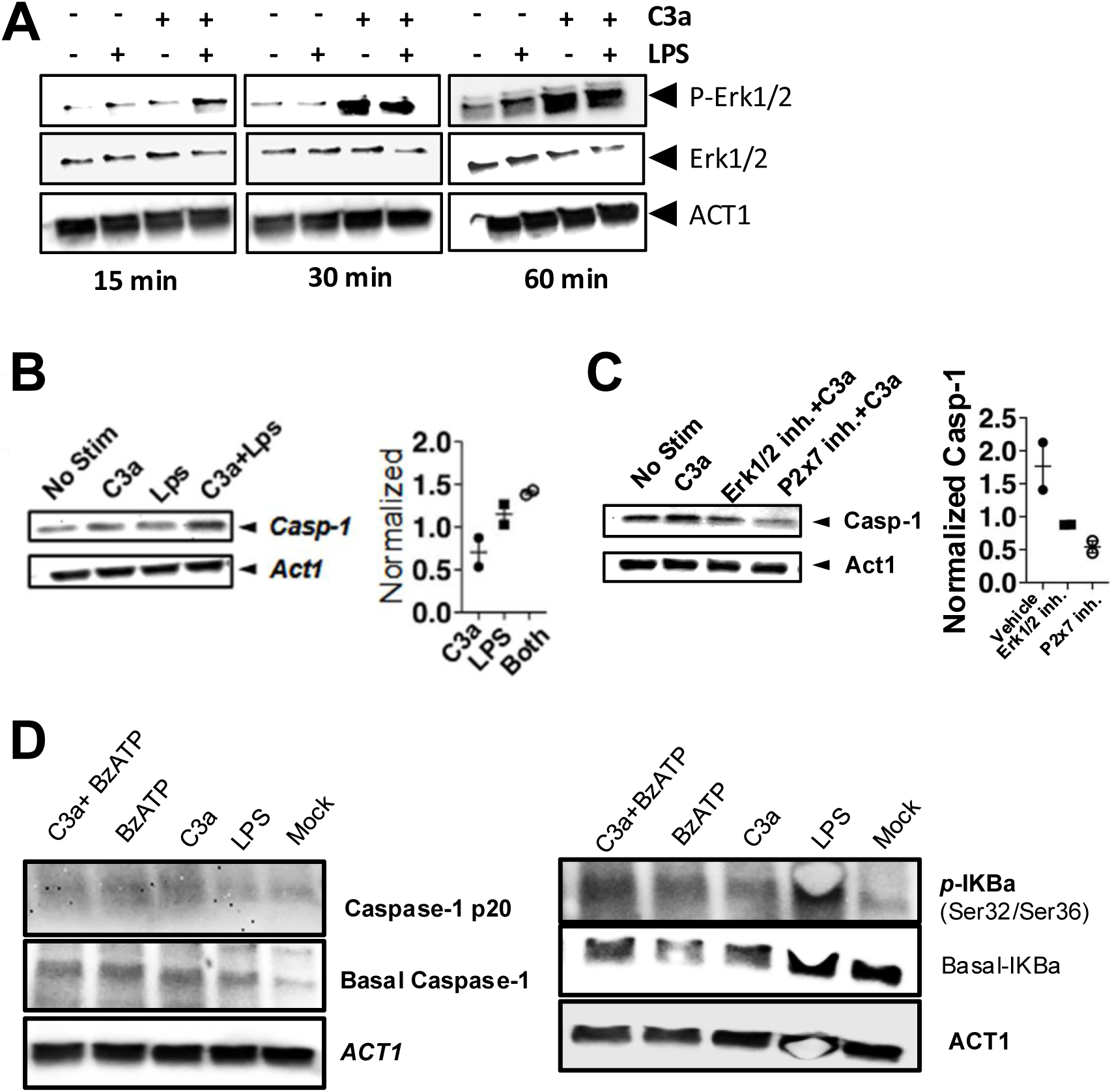
Disparate responses to C3a stimulation by distinct epithelial cell subsets. (**A**) Phosphorylation of ERK1/2 in A549 cells treated with LPS (10µg/ml), C3aR agonist (100µM) or both for 15, 30, and 60 min. (**B**) Stimulation of A549 cells with C3aR agonist and LPS induces Caspase-1 activation. Caspase-1 level in A549 cells primed with LPS (10µg/ml) and stimulated with C3aR agonist (100µM) for 24h. Densitometry quantification of Caspase-1 level normalized to mock-treated samples and *ACT1* levels. (**C**) Caspase-1 level in A549 cells after blockade of ERK1/ERK2 (10µM FR18020) or P2X7 (2µM Az11645373). Cells were primed with LPS and stimulated with C3a as above for 24h (**D**) Stimulation of SAECs with C3aR agonist induces NF-κB signaling, not caspase-1: differentiated hSAECs were apically stimulated with LPS (10 μg/ml) as a positive control, or C3aR agonist (100μM), BzATP (100 μM) or both for 12h. Cells were harvested, and extracts used to quantify of target protein level.

**Supplemental Fig. 6:**
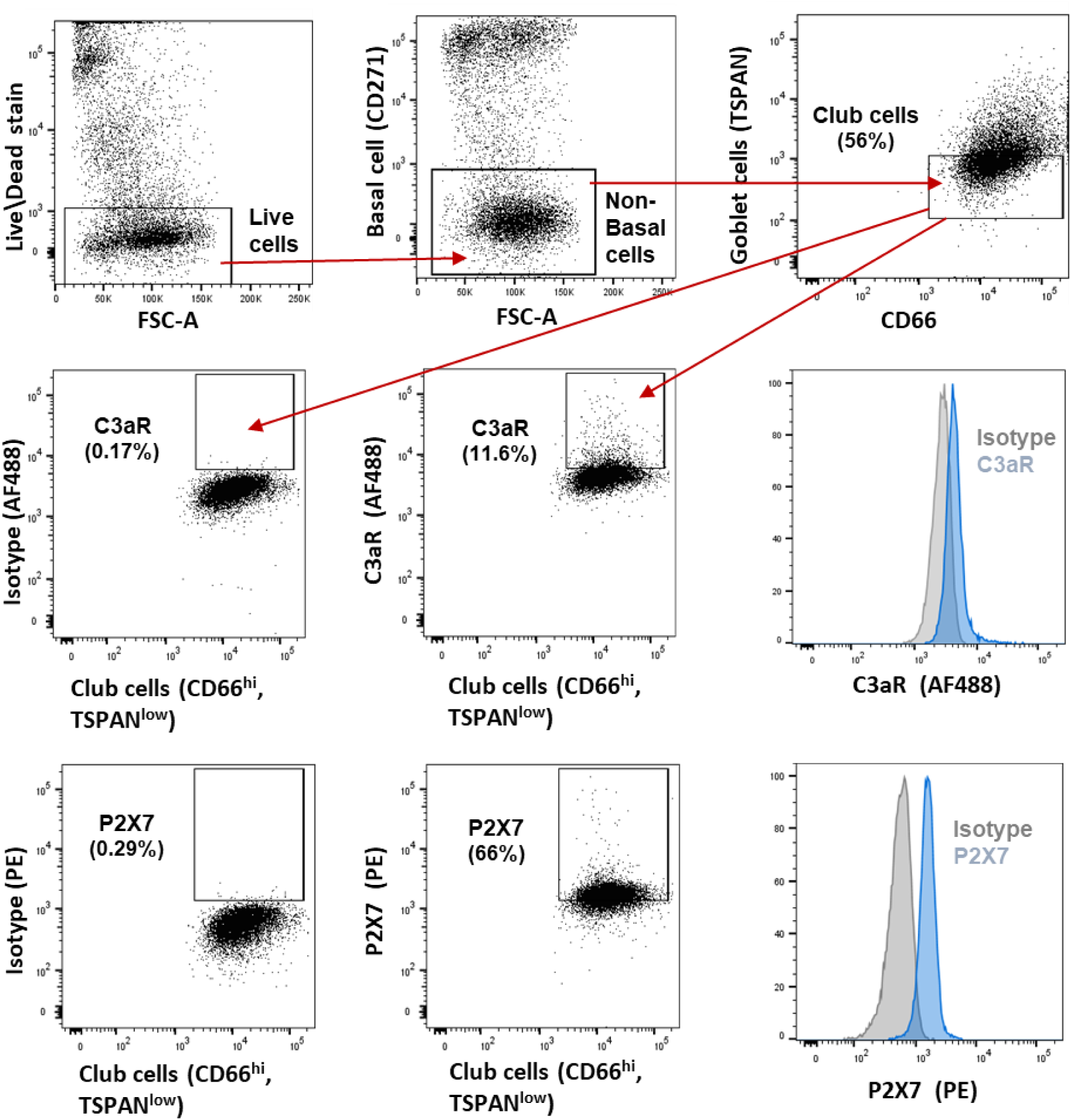
Club cells from human small airway epithelial cells express C3aR and P2X7 on the surface: Differentiated primary human airway epithelial cells were harvested after growing on specialized medium as described in Methods. Cells were removed by trypsin treatment and stained for surface markers to identify subsets with CD271 (basal cells), TSPAN (goblet cells), CD66 (club cells); and surface receptors including P2X7 and C3aR.

**Supplemental fig. 7.**
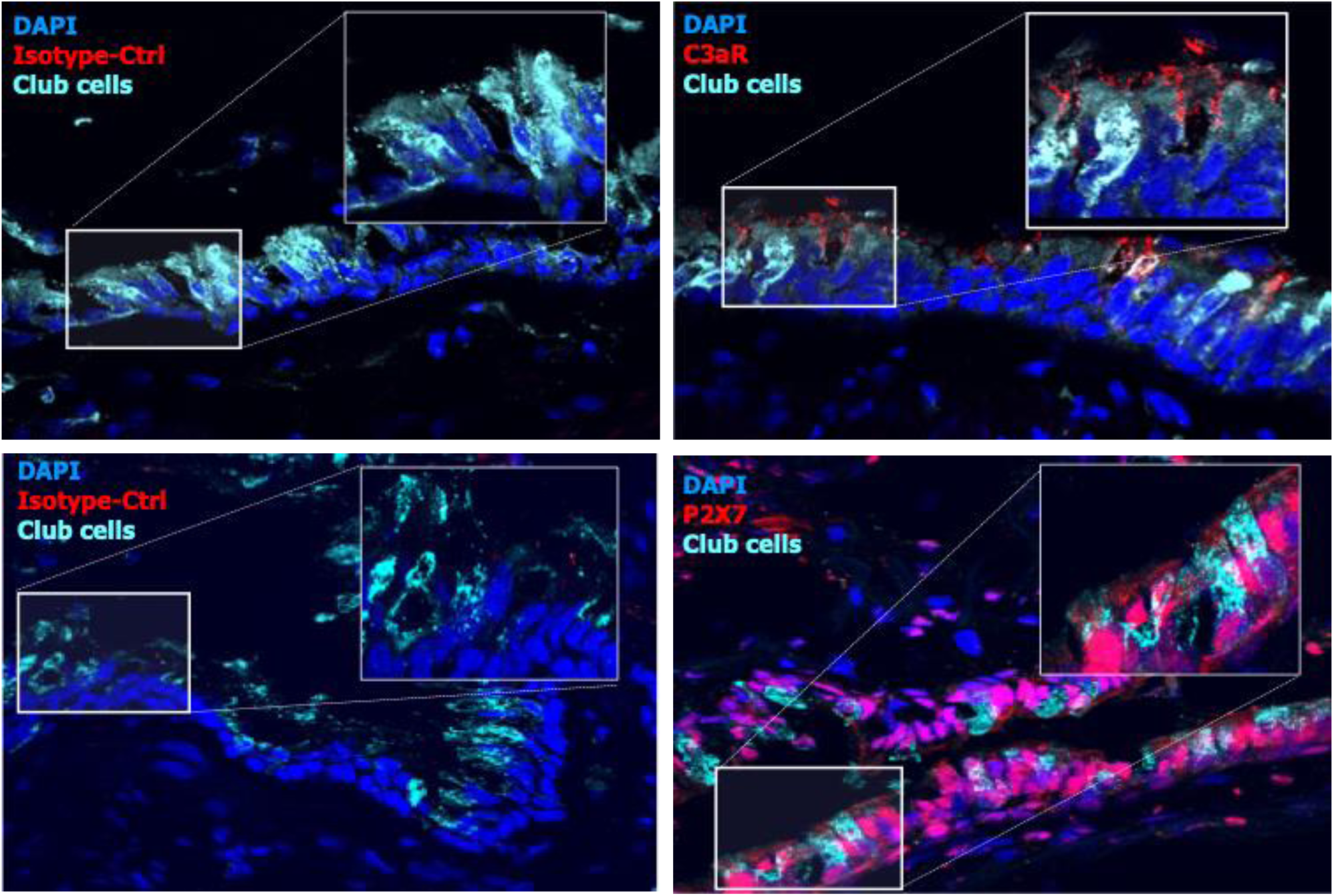
Human lung tissue staining of club cells for expression of C3aR and P2X7: Human lung sections were stained (right panels) for club cells (Scgb1A1), C3aR (hC3aRZ1) and P2X7 (epitope 576-595). Isotype control antibody staining of the receptors is depicted in left panels. Sections were stained as noted in Methods. Images were visualized at 60X on the Nikon A1R confocal microscope.

---

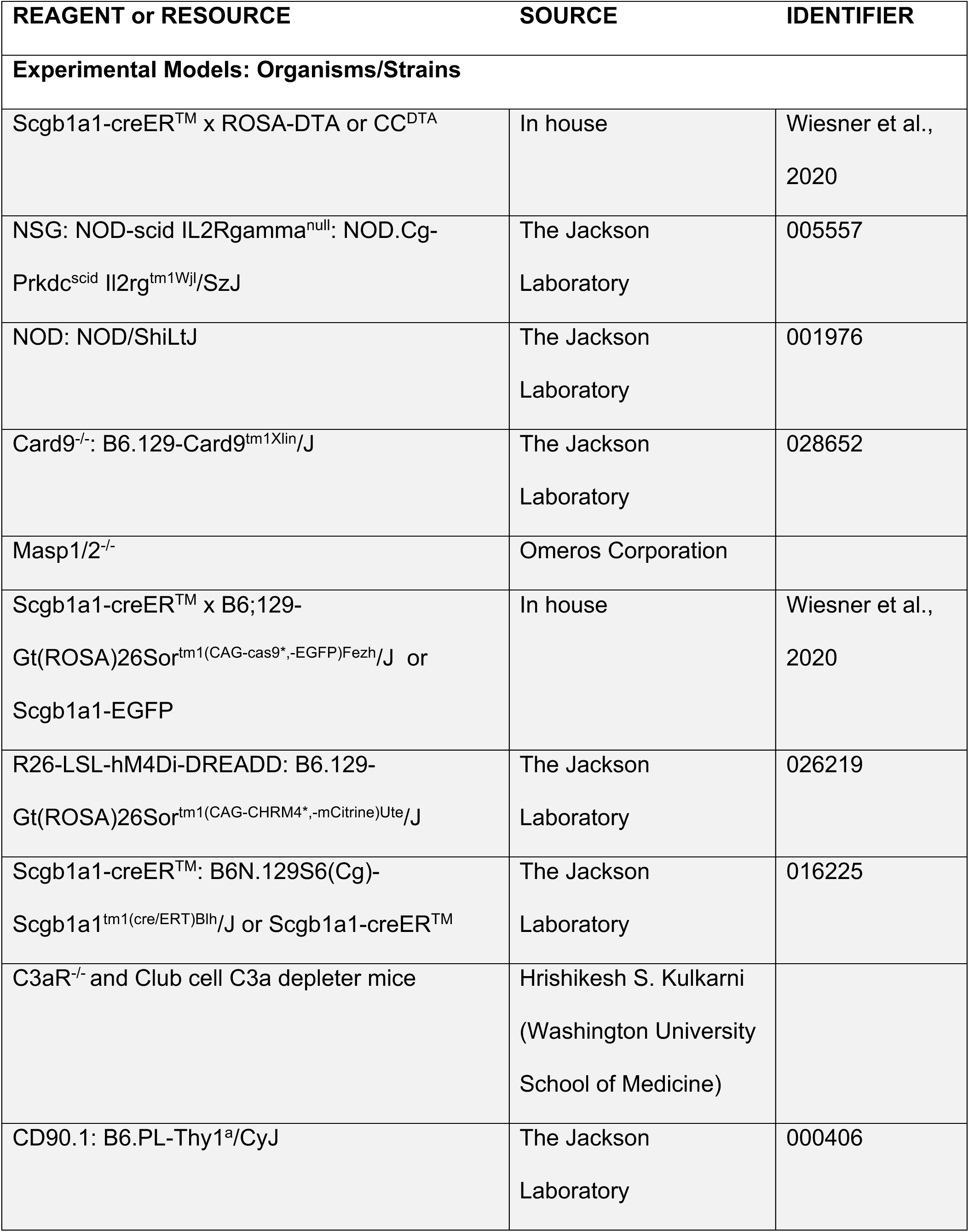
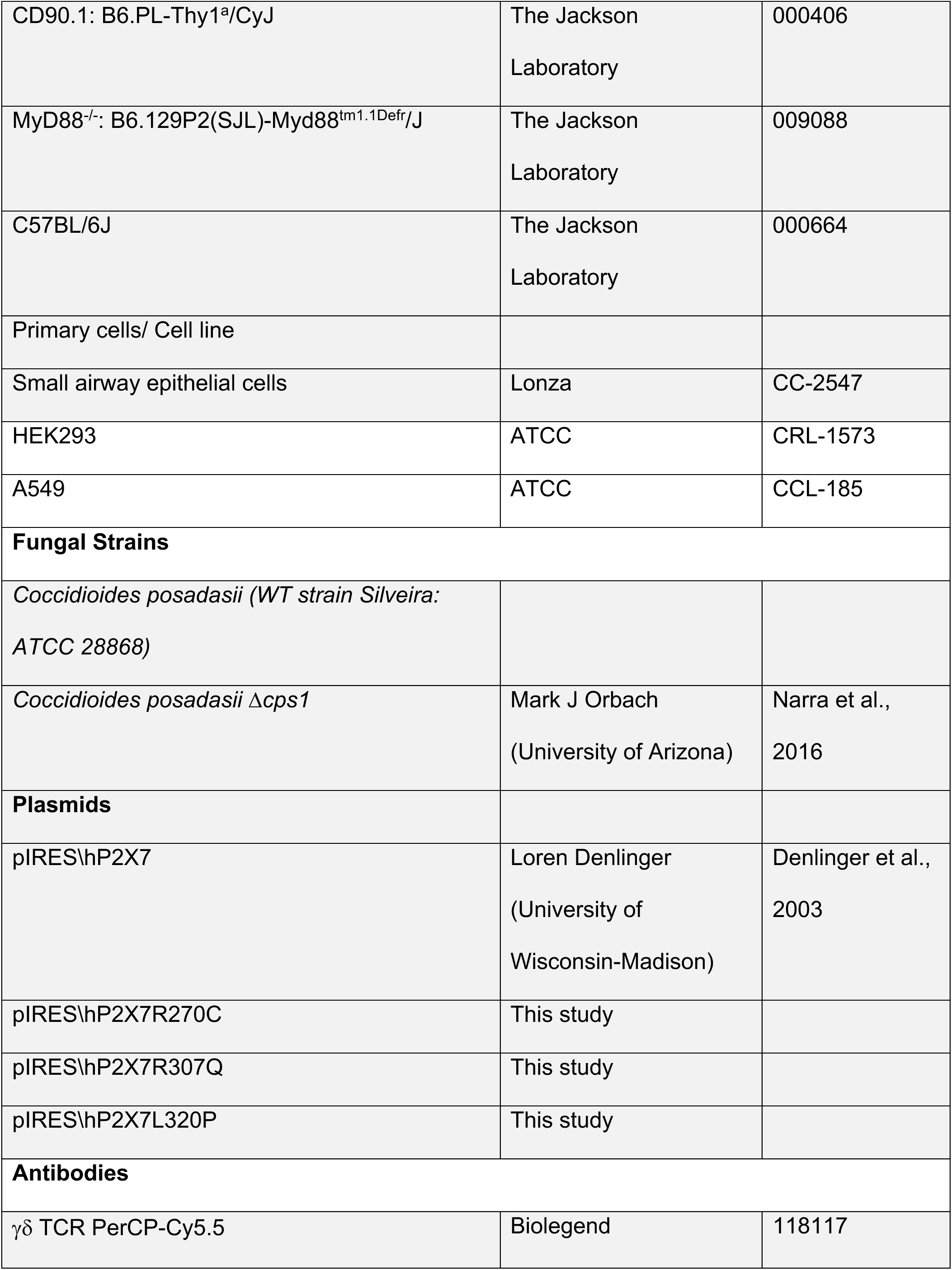
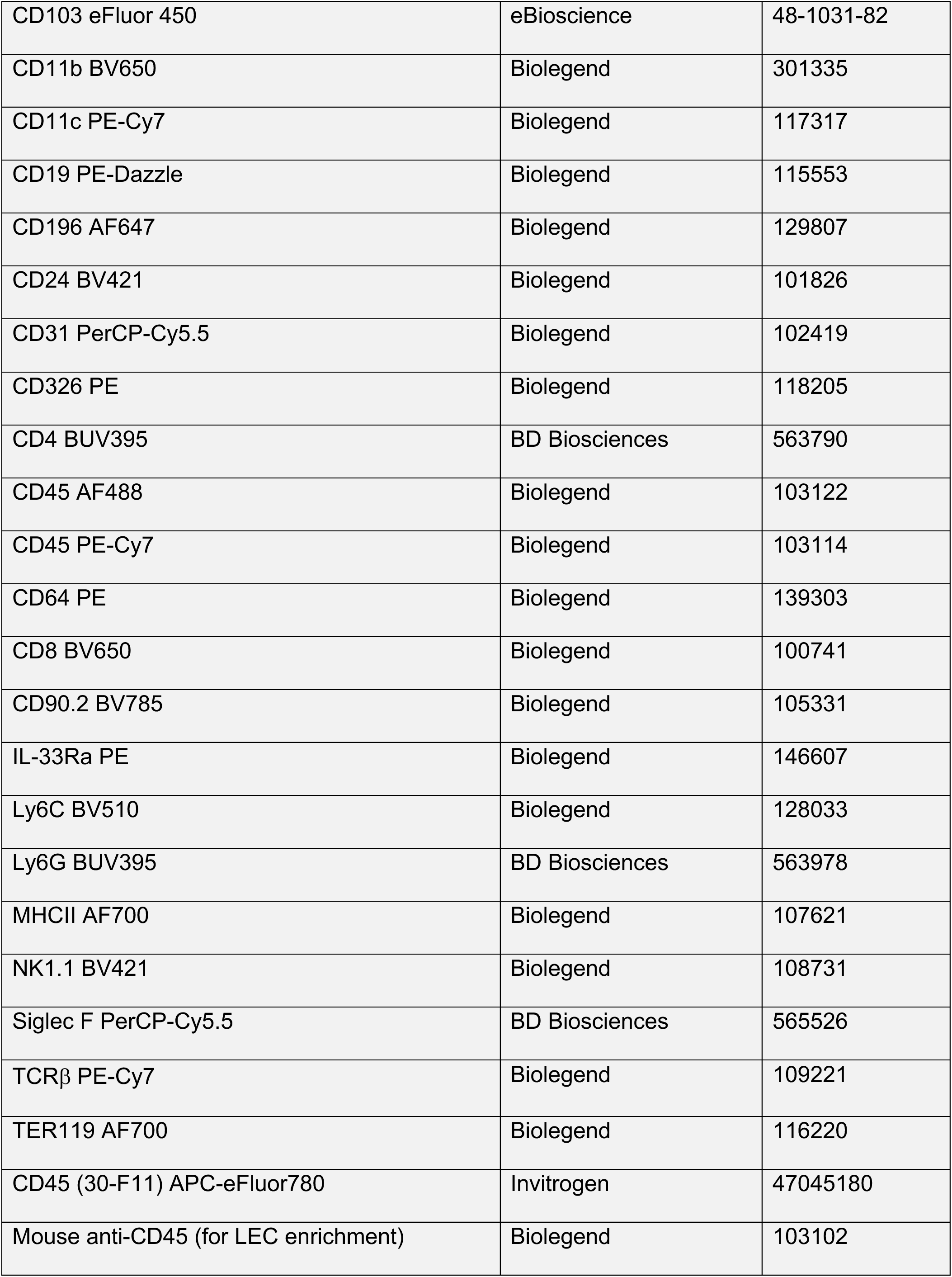
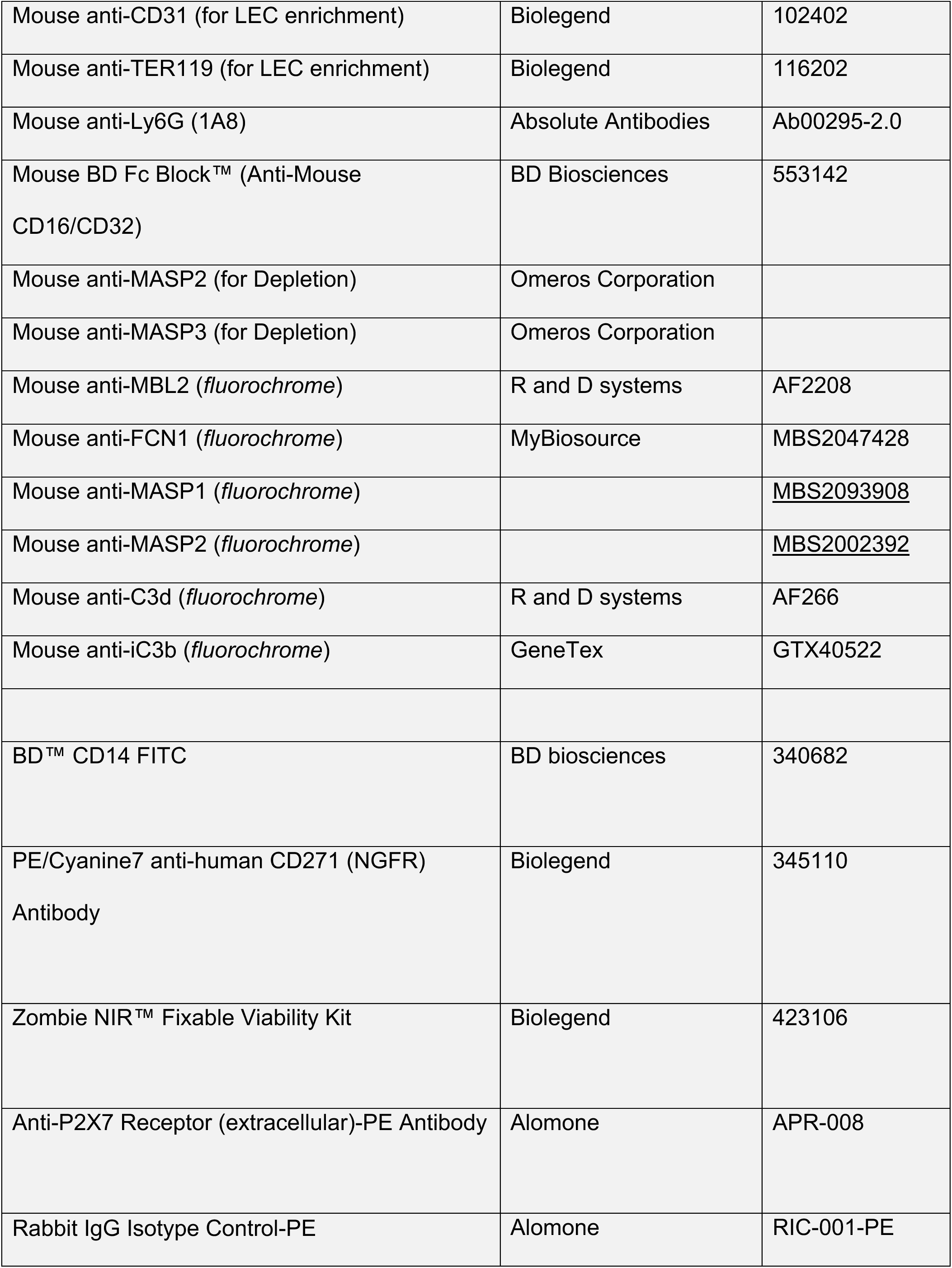
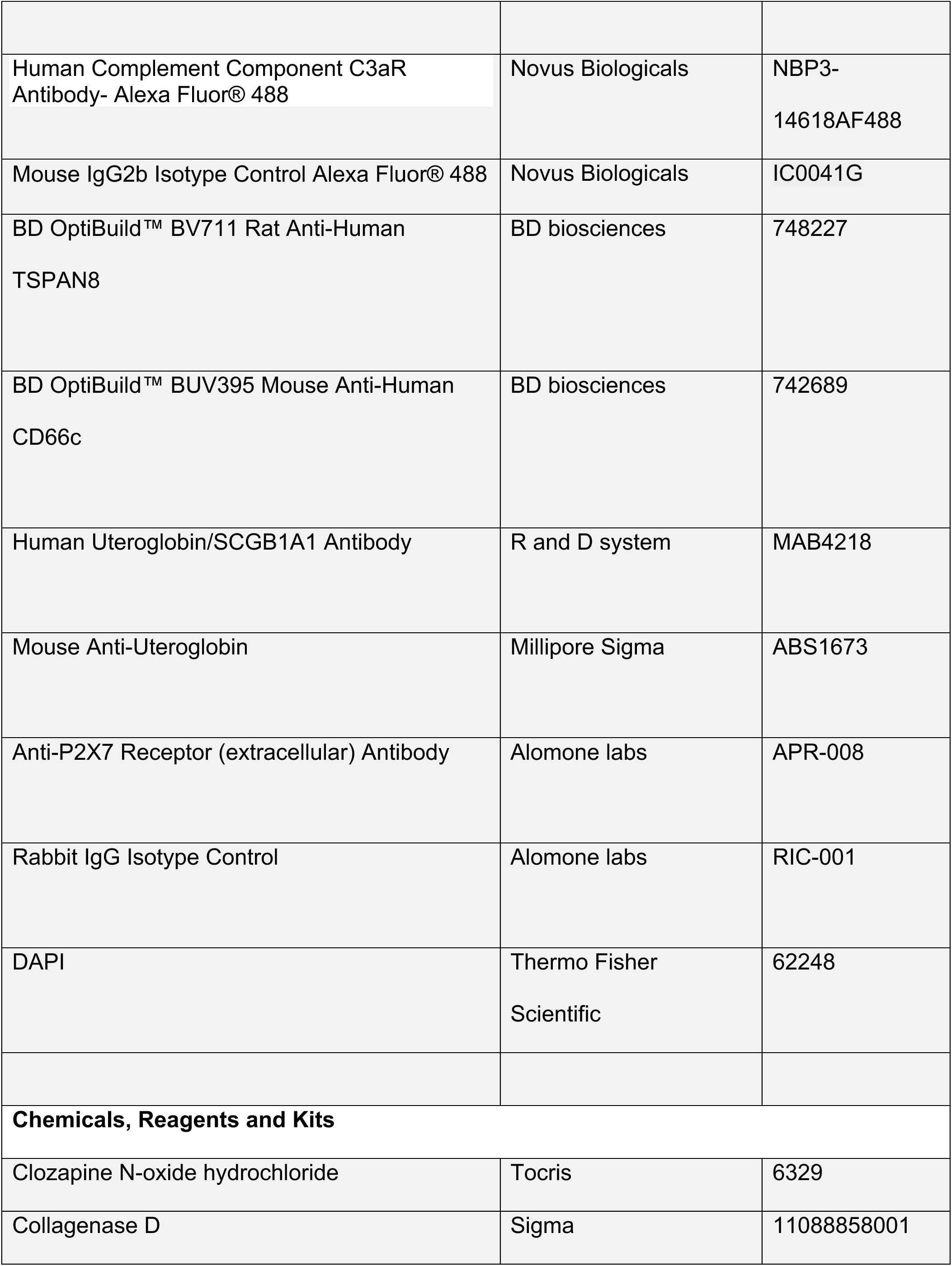
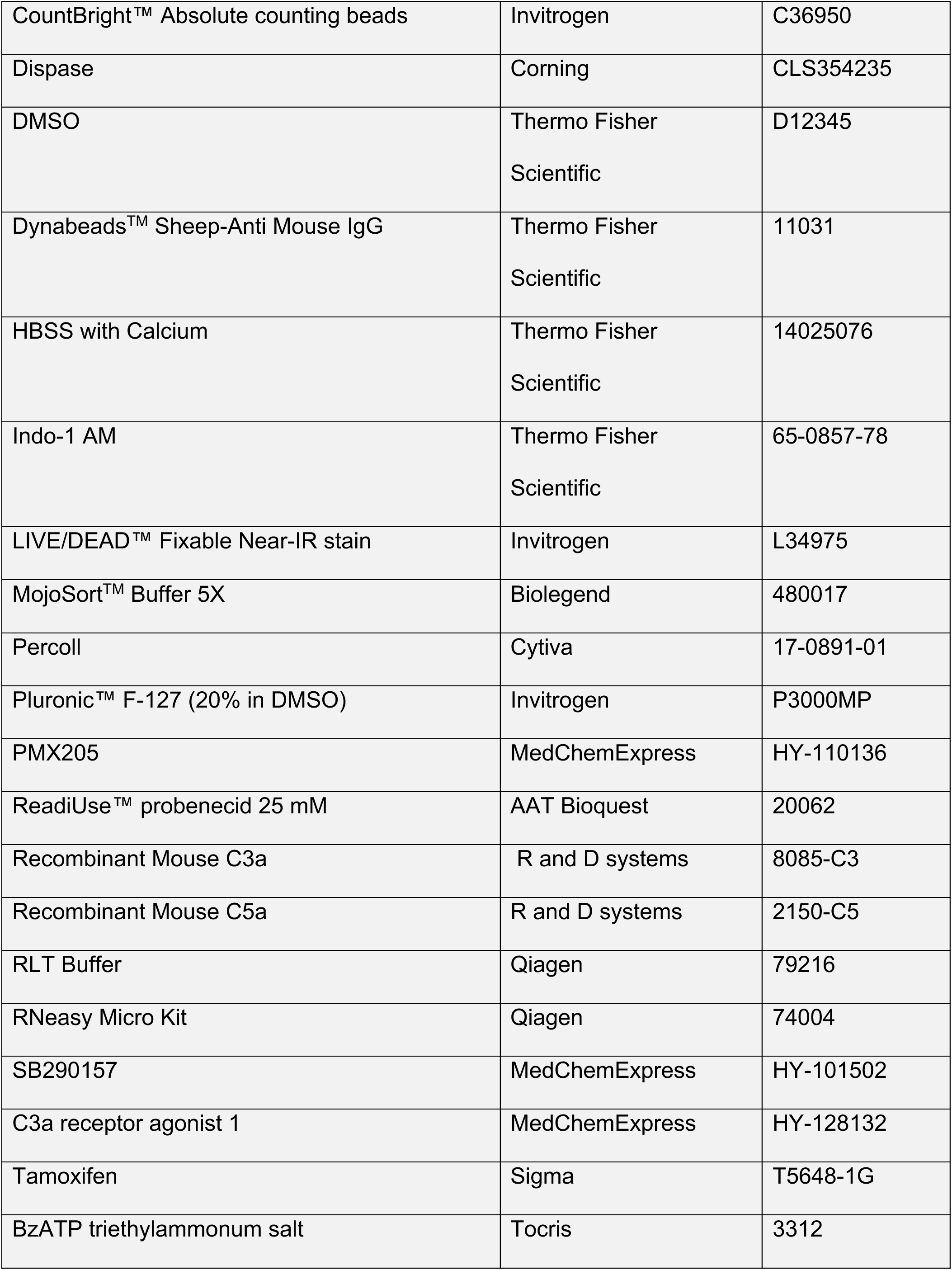
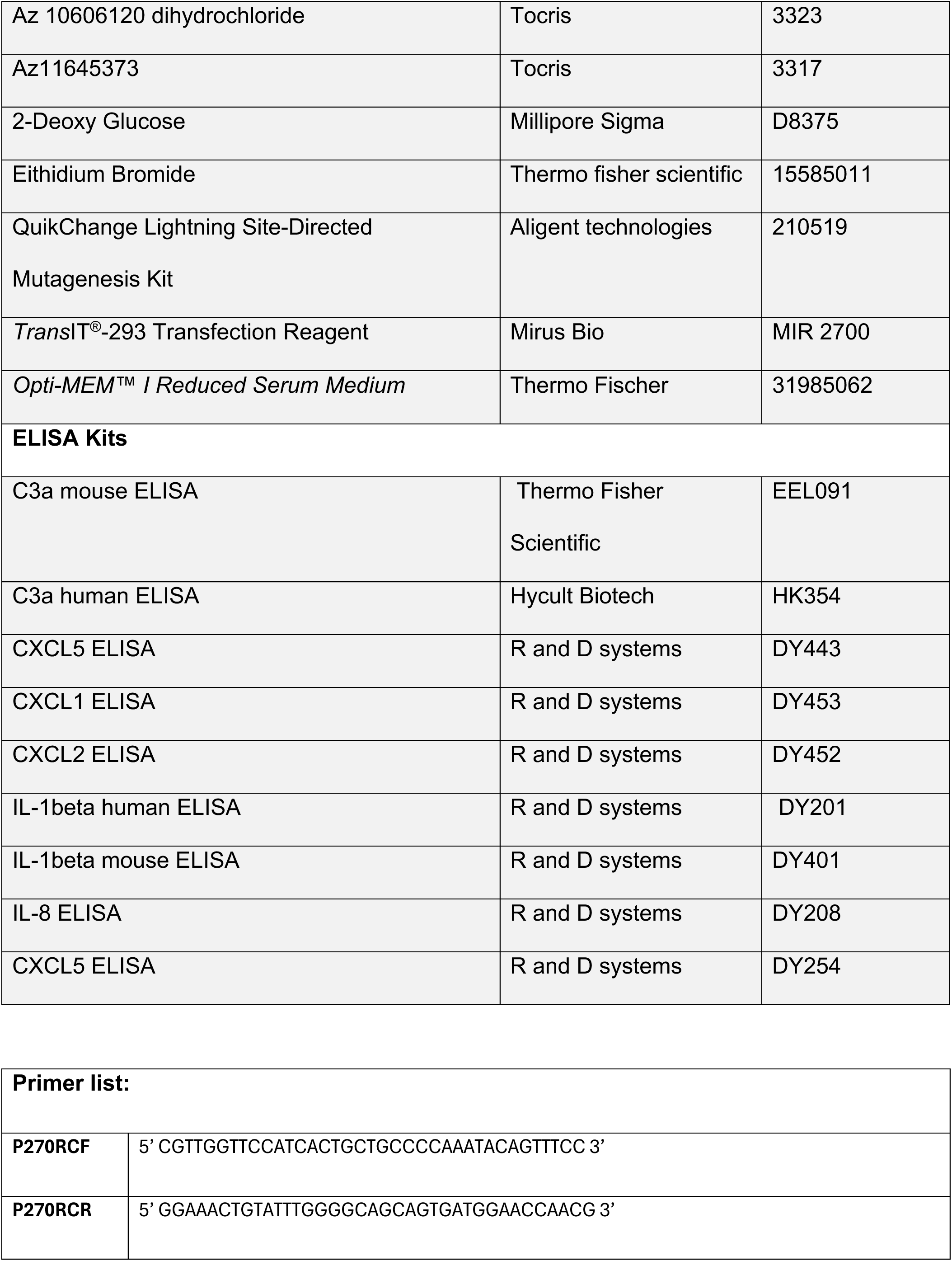
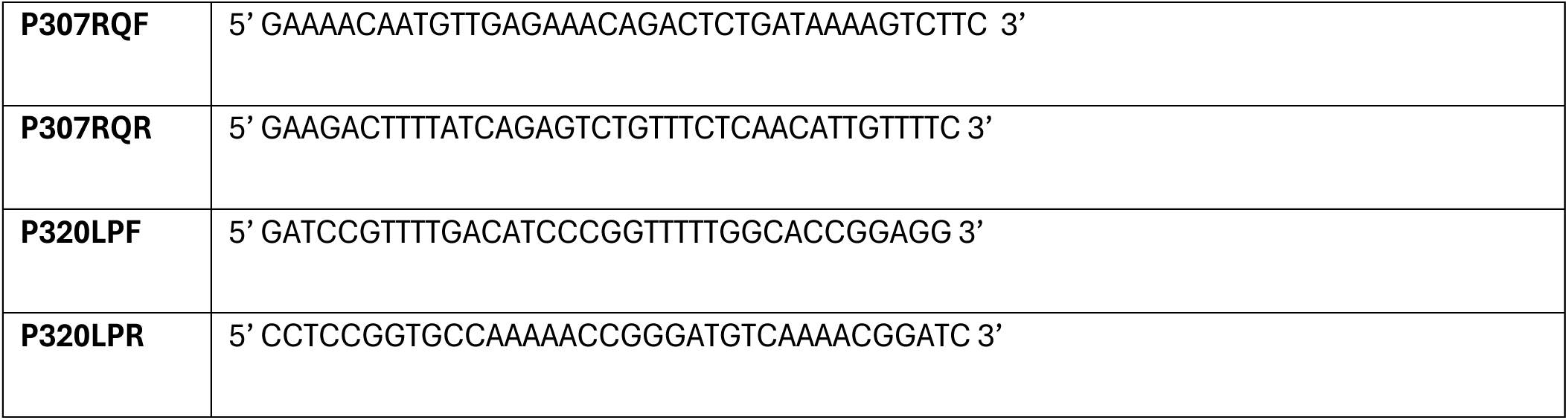
KEY RESOURCE TABLE.

## EXPERIMENTAL MODEL

Male and female mice used in study were 8 to 12 weeks of age. Wild type C57BL/6J mice and other commercial strains used were obtained from Jackson laboratory. Transgenic strains (all on a BL/6 background) included *Card9−/−, Nsg, Nod, C3ar1^−/−^* (provided by Anita Sil, UCSF), *Masp2*^−/−^^62^: club cell^DTA^ mice (Scgb1a1-creER^TM^ X ROSA-DTA) and Scgb1a1-EGFP mice (Scgb1a1-creER^TM^ X B6;129-Gt(ROSA)26Sor^tm1(CAG-cas9*,-EGFP)Fezh^/J)^63^, NSG. F1 progeny of club cell^DREADD^ mice, expressing hM4Di-DREADD in Scgb1a1 epithelial club cells, were generated by crossing Scgb1a1-creER^TM^ with R26-LSL-hM4Di-DREADD. Club cell-C3 depleter mice were generated as described^23^. Mice were bred and maintained in-house under special pathogen-free conditions and matched by sex and age in experiments for scientific rigor. Age and sex of the mice had no confounding effects in this model. Animal procedures were approved by the Institutional Animal Care and Use Committee. See Key Resources Table for details on wild type, transgenic, and knock out animals used in this study.

## METHOD DETAILS

### Coccidioides *posadasii* growth and spore production

Wild type *C. posadasii* (strain C735) or *Δcps1* were grown at 30C on 2XGYE agar plates for 3.5 weeks (Mead et al., 2020). To harvest spores, the agar surface was scrapped and the harvested mycelia transferred to a sterile conical tube containing 25-30 sterile beads and 10ml PBS. The tube was shaken gently 20 times to dispurse arthroconidia in PBS. This mixture filtered through a 70 μM strainer. The filtrate was spun at 1500rpm for 15mins at RT and washed with 20 ml PBS and resuspended in 3-5 ml PBS. Viability of *C. posadasii* C735 and *Δcps1* spores was tested by culture before infection. All work with strain C735 was performed under BSL3 conditions, and strain *Δcps1,* under BSL2 conditions.

### Mouse model of *Coccidioides posadasii* infection

Mice were challenged with either (10^2^ to 10^4^) wildtype or 1 x 10^5^ *Δcps1* mutant spores by oropharyngeal aspiration. Mice were sedated with isoflurane, and the tongue held with blunt forceps while 25 µl of spore suspension was administered into the oropharynx for lung aspiration. At various time points post-inoculation, Bronchoalveolar lavage (BAL) fluid and lungs tissue were collected for flow cytometry, cytokine quantification, colony forming unit enumeration and histology or immunohistochemistry.

For neutrophil depletion, mice were injected intraperitoneally (i.p.) with 100 µg anti-mouse anti-Ly6G 1A8 antibody 24 h before infection (Absolute Antibody)^64^. For MASP2 depletion, mice received 1 mg/kg anti-MASP2 antibody (AbD04211, Omeros Corp.) i.p. twice 18 and 2 hours before infection. All antibody depletion studies included appropriate isotype controls. For C3aR inhibition, mice were injected i.p. with 30 µg of C3aR-antagonist SB290157 twice at 12 and 2 hours before infection, and then again 12, 24, and 48 hours post infection. Control mice received saline with 3.2% DMSO. To activate Cre recombinase in cub cell^DTA^, Scgb1a1-EGFP, and club cell^DREADD^ mice, 2 mg tamoxifen was given i.p. twice, one week before and one day before infection.

### Lung immune cellular analysis

Mice were injected retro-orbitally with 3 µg of anti-mouse APC-eFluor 780-conjugated CD45 antibody for labelling cells circulating in the blood vasculature and euthanized after 3 minutes by cervical dislocation. Lungs were harvested, minced in a lung homogenizing solution (LHS) containing RPMI-1640, 5% fetal bovine serum (FBS), 0.35 mg/ml collagenase D, 1 mM CaCl_2_, 1mM MgCl_2_ and 50 µg/ml of DNAse and agitated at 37°C for 1 hour. After incubation, lungs were homogenized with a GentleMACS^TM^ dissociator (Miltenyi Biotech). The solution was filtered through a 70 µm filter, pelleted, and resuspended in 40% Percoll-RPMI medium. A 67% Percoll-PBS layer was added over the 40% Percoll-RPMI medium cell suspension, and samples were centrifuged at 2,000 rpm (650 g) for 20 min. Leukocytes at the interface were collected, washed with FACS buffer containing PBS (pH 7.4) and 0.1% FBS, resuspended in FACS buffer, and stained with 1:500 LIVE/DEAD™ fixable dye and 1:100 anti-mouse CD16/CD32 (Fc block) for 10 min. After washing with FACS buffer, the samples were incubated with 1:100 fluorochrome-conjugated primary antibodies (See Key Resource Table) for 30 min at 4°C, fixed in 10% formalin for 10 min, and analyzed by flow cytometry (LSRFortessa BD Bioscience, San Diego, CA, USA). 50,000 count beads were added for data collection. The data was analyzed with FlowJo software (FlowJo, LLC)

### *In vivo C. posadasii* spore association with leukocytes

Mice were inoculated with 1 x 10^5^ CFSE (Carboxyfluorescein Diacetate Succinimidyl Ester; 10 µM)-stained *C. posadasii Δcps1* spores. The lungs were harvested for immune cellular analysis along with recording CFSE-labelled *Δcps1* spore events by flow cytometry. Unstained *Δcps1* spores served as a negative control.

### *In vivo* complement factor binding assay on *C. posadasii* spores

Mice were inoculated with 1 x 10^5^ CFSE-stained *C. posadasii Δcps1* spores via oropharyngeal aspiration. Two hours post-challenge, BAL fluid was collected by instilling 1 ml cold PBS into the lungs and aspirating it back. The spores were isolated by centrifuging BAL fluid at 10,000 g for 10 min and incubated with 10-20 µg/ml fluorochrome-conjugated antibodies against complement factors (MBL2, MASP1, MASP2, FCN1, iC3b, or C3d) in PBS (pH 7.4) for 30 mins at 4°C. After incubation, spores were washed thrice, fixed with 1% formaldehyde in PBS, and quantification of complement factors binding was done via flow cytometry, with negative controls being unchallenged CFSE-stained *Δcps1* spores.

### C3a detection by ELISA

150,000 to 200,000 *Δ*cps1 spores resuspended in 50ml of PBS and incubated with 50ml human plasma for 0h, 2h, 4h and 6h. Samples were spun down at 1500 rpm for 10 min at 4^◦^C. Supernatant was collected and C3a levels were quantified by ELISA (Hycult Biotech).

### Lung epithelial cells (LEC) enrichment and calcium flux

Tamoxifen-treated Scgb1a1-creER^TM^ x LSL-Cas9 GFP mice (or Scgb1a1-EGFP)^63^ were euthanized by CO_2_ asphyxiation, and lungs were instilled with 1 ml of Dispase (50 Units/ml) via the trachea. The trachea was sealed with a thread and the lungs were incubated in LHS at 37°C for 30 min at 100 rpm. After incubation, lungs were homogenized and the cells suspension was filtered through 100 µm and 40 µm strainers. Erythrocytes were lysed with Ammonium Chloride Potassium lysis buffer. The cells were washed with FACS buffer, pelleted and resuspended in 1X MojoSort^TM^ buffer containing 2 mM EDTA for enrichment of lung epithelial cells via magnetic depletion of antibody bound unwanted cells in samples. The samples containing 1 x 10^7^ cells were labeled with a cocktail of antibodies (10 µg each of Rat IgG anti-mouse CD45, CD31 and TER119) at 4°C for 20 min, followed by incubation with magnetic Dynabeads^TM^ sheep-anti mouse IgG at 4°C for 30 minutes. Post-incubation, samples were magnetically separated to capture Dynabeads-bound non-epithelial cells, allowing lung epithelial cells to remain in the supernatant. The enriched epithelial cells suspension was collected, loaded with 5 µM Indo-1 AM with 0.02% pluronic acid, 4 mM ReadiUse probenecid, and 10 mM HEPES (pH 7.4) for 30 min at 37°C, then washed with PBS without calcium and magnesium, 10 mM HEPES (pH 7.4) and 1% FBS buffer. The Indo-1 loaded cells were stained with fluorochrome antibodies, resuspended in HBSS with 1 mM calcium, 10 mM HEPES (pH 7.4) and 1% FBS buffer and analyzed by flow cytometry. Baseline fluorescence for Indo-1 was recorded at dual wavelengths of 475 nm (calcium-free) and 400 nm (calcium-bound) for 20 seconds before adding 50 µl bronchoalveolar lavage fluid collected from the lungs of *Δcps1* spore-challenged mice to induce calcium flux. Recombinant mouse C3a or C5a protein were used as positive controls for recording the activation of C3a– or C5a-receptor mediated calcium flux, respectively. To inhibit C3a– or C5a-receptor activity, Indo-1 loaded cells were incubated with 10 µM SB290157 or 10 µM PMX205, respectively, for 10 minutes at 37°C prior to data collection.

### Bone marrow chimera experiments

Two doses each of 550 Gray lethal irradiation were administered to recipient mice (C57BL/6 or C3aR^−/−^) at 3-hour intervals. Post-irradiation, mice were kept in a sterile environment. 24 h later, the mice received 1 x 10^6^ donor bone marrow (C57BL/6J or C3aR^−/−^) cells retro-orbitally. Two months later, the chimeric mice were challenged with *Δcps1* spores and the lungs were harvested for immune cellular analysis by flow cytometry.

### Epithelial Cell sorting and RNA preparation

To obtain EpCAM+ (CD326+) CD24+ airway club cells for RNA preparation, mice were euthanized by CO_2_ asphyxiation, and lungs were collected. The lungs were homogenized with Dispase treatment and GentleMACS^TM^, and the resulting cell suspension was filtered through a 40 µm strainer. The single-cell suspension was stained with 1:500 LIVE/DEAD^TM^ fixable dye and 1:100 Fc block, followed by fluorescent antibodies for leukocytes (anti-CD45), erythrocytes (anti-TER119), endothelial cells (anti-CD31), epithelial cells (anti-EpCAM), and club cells (anti-CD326, anti-CD24). The cells were washed with FACS buffer, pelleted and resuspended for flow cytometry and sorting. A total of 500,000 CD45-CD31-TER119-CD326+CD24+ club cells were collected in 1 ml of RLT RNA lysis buffer containing β-mercaptoethanol (Qiagen). The sorted cells were vortexed and snap-frozen at –80°C for RNA isolation, which was performed using the RNeasy Micro Kit following the manufacturer’s instructions with two additional washes.

### RNA Sequencing, Enrichment, and Visualization

RNA quality and concentration was confirmed using an Agilent Bioanalyzer, with RIN values ranging from 8.5-9.5. Libraries were generated using the Takara SMARTer Stranded Total RNA-Seq Kit v2– Pico Input Mammalian kit with 10ng of total RNA input, 4 minutes of fragmentation, and 12 PCR cycles of amplification, generating libraries with fragment sizes ranging from 367 to 436 base pairs. Libraries were sequenced on an Illumina sequencer at a target depth of 40-45 million mapped reads per sample. Sequencing quality control metrics were obtained using FASTQC Multiqc^65^ and sample reads were aligned to the mm10 genome using STAR^66^ and quantified using featureCounts^67^. Differential expression testing was performed using EdgeR glmqlfit^68^. Gene ontology analysis was performed on DEG with FDR <0.05 and absolute log-fold change >0.6 using Metascape^69^. Heatmaps were generated with the ComplexHeatmap^70^ package in R.

### Chemogenetic analysis of the role of GPCR on Club Cells

Scgb1a1-creER^TM^ mice were crossed with R26-LSL-hM4Di DREADD mice to generate CC^DREADD^ mice, which express a mutant G protein-coupled receptor, hM4Di-DREADD, in Scgb1a1 expressing lung epithelium club cells. The hM4Di DREADD activates the G_i_-inhibitory signaling pathway after administration of pharmacologically inert molecule clozapine-N-oxide (CNO), inhibiting adenylyl cyclase and lowering cAMP levels. For hM4Di-DREADD induction, club cell^DREADD^ mice received 2 mg tamoxifen i.p. twice, 7 days and 1 day before infection. 3 mg/kg CNO was given 1 hour before challenge with *C. posadasii* spores and every 12 hours later. Lungs were harvested and analyzed for immune cells. Controls included C57BL/6 mice treated with tamoxifen plus CNO and club cell^DREADD^ mice treated with tamoxifen alone.

### P2X7 function assayed by Ethidium Bromide (EtBr) uptake in peripheral blood mononuclear cells (PBMC)

As described previously^31^. frozen PBMCs were thawed and 1ml of complete RPMI (with 10% FBS and 1% P/S) was added drop wise to the vial. Thawed cells were added to additional RPMI, incubated at 37^◦^C for 5min, centrifuged at 330 g for 10min and resuspended in 1ml of complete RPMI medium with DNase-I (50 Units/ml). After cells were incubated for 1hr at 37^◦^C, viability was quantified with trypan blue. Viable cells were transferred into a 96-well U-bottom plate (10^6^/well) and washed twice with NaCl medium (145mM NaCl, 5mM KCl, 5mM glucose, 10mM HEPES pH 7.4. Cells were treated with Fc Block Ab for 5min, washed once with NaCl medium and resuspended in 100μl of NaCl medium. 2μl of BzATP, 2μl of CD14 Ab (conjugated with FITC) and 1μl of EtBr (2.5mM) were added to each well and incubated for 10-12 min at 37^◦^C. To stop EtBr uptake by PBMC, 100μl of cold MgCl_2_ buffer (20mM MgCl_2_, 145mM NaCl, 5mM KCl, 5mM glucose, 10mM HEPES, pH 7) was added to each well. Cells were washed, resuspended in 150μl of NaCl medium, and placed on ice until the samples were analyzed on a BD Fortessa.

### Construction of P2X7 variants

The pIRES-P2X7 wild-type construct^71^ was used to generate the R270C, R307Q, and L320P P2X7 variants by site-directed mutagenesis with the QuikChange Lightning kit (Agilent). Mutagenesis was performed using mutation-specific primers following the manufacturer’s protocol. After PCR amplification, parental methylated plasmid DNA was digested with DpnI, and the resulting products were transformed into DH5α competent bacteria. Colonies were selected on LB-ampicillin plates, expanded, and plasmid DNA was purified for sequence verification. All constructs were confirmed by Sanger sequencing.

### HEK 293 transfection and EtBr uptake assay

HEK293 cells (1×10⁵) were plated in a 6-well plate in DMEM containing 10% FBS (without antibiotics) and allowed to adhere for 16 h. Cells were transfected using the TransIT-293 reagent according to the manufacturer’s protocol. 2.5 µg plasmid DNA was mixed with 7.5 µL TransIT-293 reagent in 250 µL Opti-MEM I reduced-serum medium and added to each well. After 6 h, the medium was replaced with fresh DMEM containing 10% FBS (without antibiotics). 48hrs after transfection, cells were washed with 1x PBS, harvested, and counted using a hemocytometer. HEK293 cells expressing wild-type P2X7 or the indicated variants were resuspended and 1 × 10⁶ cells were transferred to a 96-well U-bottom plate and washed twice with NaCl medium (described above). Cells were stained with Zombie NIR viability dye for 5 min and washed once with NaCl medium. EtBr uptake was initiated by adding 1 µL EtBr (2.5 mM) and 2 µL BzATP to each well, followed by incubation at room temperature for 8 min. Uptake was stopped by adding 100 µL ice-cold MgCl₂ buffer (described above). Cells were washed, resuspended in 150 µL NaCl medium, and kept on ice until analysis on a BD Fortessa flow cytometer.

### Cell signaling

A549 cells were grown in RPMI medium (10% FBS +1% P/S). Cells were starved in serum free medium overnight and stimulated with C3a agonist (75μM) for 15 to 60min. For LPS stimulated samples, cells were primed with LPS for 4 hours with 10μg/ml, followed by C3a stimulation (100μM). For inhibitor studies, cells were treated with Erk1/2 inhibitor (FR 180204) 5 μM for 30min or P2X7 inhibitor (AZ 10606120) 2.5μM for 30min. For western blotting, after stimulation cells were washed with 1X PBS and scrapped off the surface with 1X RIPA buffer containing protease and phosphatase inhibitor. 20-30µg of protein extract was loaded on the SDS PAGE followed by immunoblotting to detect the target protein.

### Primary small airway epithelial cells

Small airway epithelial cells (Cat. No. CC-2547) were acquired from Lonza Integrated Biologics. Cells were grown in PneumaCult™-Ex Plus Medium basal medium for one week, and differentiation medium PneumaCult™-ALI-S Medium (Cat. No. 05050) for 3 weeks in a transwell plate. Differentiated cells were treated apically with C3a (100µM). Post stimulation, samples were collected from the apical surface by adding 100-200μl of PBS.

### P2X7 antagonist and 2-DG

P2X7 antagonist (AZ 10606120) was given IP (2mg/kg), on days –2, –1 and 0 before infection with *Δcps1* spores. 2-Deoxy Glucose (Sigma-Millipore Cat. No. D8375) was dissolved in sterile water was given (1mg/kg) on days –2, –1 and 0, and again daily until the experiment was terminated.

### Immunohistochemistry (IHC) of human or mouse lung tissue

Human lung tissue blocks were acquired from NDRI and sectioned on a frozen microtome with 10-micron thickness. Tissue sections on slides were fixed with 10% formalin at room temperature for 20min, followed by 3 washes with PBS. Lung tissue sections were stained as described^63^.

### Quantification And Statistical Analysis

Statistical analyses comparing more than two groups were Mann-Whitney U test with Bonferroni adjustment for multiple comparisons using Graph-Pad Prism 9 (San Diego, CA). A p value <0.05 was deemed statistically significant, and error bars are SEM. All data represent at least two independent experiments.

### Genetic association

*Genomic boundaries for variants and variant nomenclature.* Variants falling within the following hg19 regions were extracted from the variant callset files of the validation cohort and 1000G: *P2RX7* – NM_002562.6, chr12:121132819-121188032; *MBL2* – NM_001378373.1, chr10: 52765380-52772784; *C3AR1* – NM_004054.4, chr12:8,056,844-8,066,359. Variants were annotated using CADD version 1.7 (Schubach M, Maass T, Nazaretyan L, Roner S, Kircher M, PMID:38183205)

*Case–control matching.* PCA of the genotype data was performed using ∼10,000 ancestry-informative variants, after which 4 population controls from the 1000G (*n* = 2504) were selected for each DCM case using the R package *Optmatch* (https://cran.r-project.org/web/packages/optmatch/index.html), based on PC1–PC5.

*Statistical analysis of variant burden.* A logistic regression model for the binary response of disseminated disease status was used to estimate the OR of variant burden in our genes of interest, adjusting for principal components of ancestry background as previously reported. Three gene association tests (*P2RX7, MBL2*, and *C3AR1*) were performed. Predicted damaging variants were selected using gnomAD_controls_AF_popmax <0.1, CADD (version 1.7) phred score ≥20, and “PASS” filter status in the variant callset file. A *P* value < 0.05 (2 tailed) was considered significant and a gene OR of greater (less) than 1 is interpreted as indicating positive (negative) association. Bonferroni adjustment was used to correct for multiple testing. Association tests were then performed for recurring variants.

